# Adrenergic signaling coordinates distant and local responses to amputation in axolotl

**DOI:** 10.1101/2021.12.29.474455

**Authors:** Duygu Payzin-Dogru, Tim Froitzheim, Steven J. Blair, Siddhartha G. Jena, Hani Singer, Julia C. Paoli, Ryan T. Kim, Emil Kriukov, Sarah E. Wilson, Renzhi Hou, Aaron M. Savage, Victor Cat, Louis V. Cammarata, S. Y. Celeste Wu, Vivien Bothe, Burcu Erdogan, Shifa Hossain, Noah Lopez, Julia Losner, Juan Velazquez Matos, Sangwon Min, Sebastian Böhm, Anthony E. Striker, Kelly E. Dooling, Adam H. Freedman, Bobby Groves, Benjamin Tajer, Glory Kalu, Eric Wynn, Alan Y. L. Wong, Nadia Fröbisch, Petr Baranov, Maksim V. Plikus, Jason D. Buenrostro, Brian J. Haas, Isaac M. Chiu, Timothy B. Sackton, Jessica L. Whited

## Abstract

Many species regenerate lost body parts following amputation. Most limb regeneration research has focused on the immediate injury site. Meanwhile, body-wide injury responses remain largely unexplored but may be critical for regeneration. Here, we discovered a role for the sympathetic nervous system in stimulating a body-wide stem cell activation response to amputation that drives enhanced limb regeneration in axolotls. This response is mediated by adrenergic signaling, which coordinates distant cellular activation responses via the α_2A_-adrenergic receptor, and local regeneration responses via β-adrenergic receptors. Both α_2A_– and β-adrenergic signaling act upstream of mTOR signaling. Notably, systemically-activated axolotls regenerate limbs faster than naïve animals, suggesting a potential selective advantage in environments where injury from cannibalism or predation is common. This work challenges the predominant view that cellular responses underlying regeneration are confined to the injury site and argues instead for body-wide cellular priming as a foundational step that enables localized tissue regrowth.

## INTRODUCTION

The ability to spontaneously replace large body parts lost to injury or disease varies across the animal kingdom. Salamanders, such as axolotls, for example, can regenerate full limbs, but mammals cannot. Relatively little is known about how selective pressures experienced by salamanders might have shaped their remarkable regenerative traits. Furthermore, despite progress in understanding the molecular responses of salamander cells at the amputation site, how cells elsewhere in the body respond to amputation of a distant limb and the implications of these responses have been less explored. Yet, these distant responses could be important for understanding limb regeneration and the evolution of this trait, and they could provide essential insights for future regenerative medicine approaches aimed at harnessing latent regenerative capacity in humans.

Previously, we found that in axolotls, limb amputation provokes a body-wide cellular proliferation response within many distant tissues in a process we called “systemic activation.”^1^ Systemic activation occurs even when limbs are experimentally blocked from regenerating, indicating that it is not likely to be driven by growth factors secreted by regenerating limbs. However, the molecular signals that stimulate systemic activation remained unknown. In this earlier work, we also found that an ancient pathway controlling cell growth across many species, mTOR, is enhanced both at the amputation site and within distant, responding tissues. However, whether mTOR signaling is required for systemic activation in axolotls, as well as the potential upstream signals that might regulate mTOR in this context, remained unknown. A further mystery has been whether the signaling pathways that govern systemic activation are unique to distant locations, or whether they might also drive cellular proliferation and blastema formation at the amputation site. We therefore sought to uncover how distant cells are cued by amputation to become systemically activated in axolotl, the relationship of these stimuli to mTOR signaling, and their potential to promote blastema formation at the injury site. In this work, we also aimed to uncover the implications of systemic activation on axolotls.

Here, we show that systemic activation primes distant limbs for faster regeneration in axolotls, a response that may be evolutionarily relevant in wild salamander populations. We demonstrate that peripheral innervation of both the amputation site and the distant, responding site is required for systemic activation. We further find that the sympathetic nervous system mediates this dependency through α-adrenergic signaling. We also show that mTOR signaling operates downstream of α-adrenergic signaling in this process. Interestingly, while α-adrenergic signaling is required for systemic activation, β-adrenergic signaling is dispensable for this process. However, β-adrenergic signaling is necessary at the amputation site for limb regeneration, where it promotes mTOR signaling, cellular proliferation, and blastema formation. Our work demonstrates that a common signal, the adrenergic ligand norepinephrine, is interpreted by cells differently depending on their proximity to the amputation site. In the amputated limb tissues, both α_2A_– and β-adrenergic receptor activities promote mTOR signaling and proliferation by blastemal progenitors, while at distant sites elsewhere in the body, α-adrenergic receptors stimulate mTOR to promote systemic activation, thereby priming axolotls for enhanced regenerative responses to future injuries. These results highlight potentially evolutionarily relevant responses to amputation that could be explored in other species to promote regenerative therapies.

## RESULTS

### Amputation primes distant uninjured limbs for future local regeneration

Recently, we uncovered a phenomenon of body-wide cellular proliferation response to amputation in axolotl (*Ambystoma mexicanum*). Here, we sought to understand the consequences of systemic activation to axolotls. We first asked whether systemically-activated axolotls are more, or less, effective at mounting future regenerative responses. We challenged systemically-activated axolotls with amputation (**Figure 1A**) and found they regenerate these limbs, which are distant to the originally-amputated limbs, significantly faster than naïve control animals (**Figure 1B**). Systemically-activated limbs grew blastemas more quickly (**Figure 1C**) and reached digit-differentiation stages faster (**Figure 1D-E**). We asked whether this enhanced regeneration response is facilitated by more rapid blastema cell specification and coalescence. We visualized the expression of an established blastema transcript, *Prrx1*, using HCR, and found that systemically-activated limbs have more robust blastema-initiation responses than naïve limbs (**Figure 1F-G**). This data supports the conclusion that amputation-induced systemic activation leads to priming of distant tissues for faster regeneration in animals subjected to future local injury. A faster recovery of digits, for example, may be critical for animal fitness and survival in the wild.

**Figure 1.**
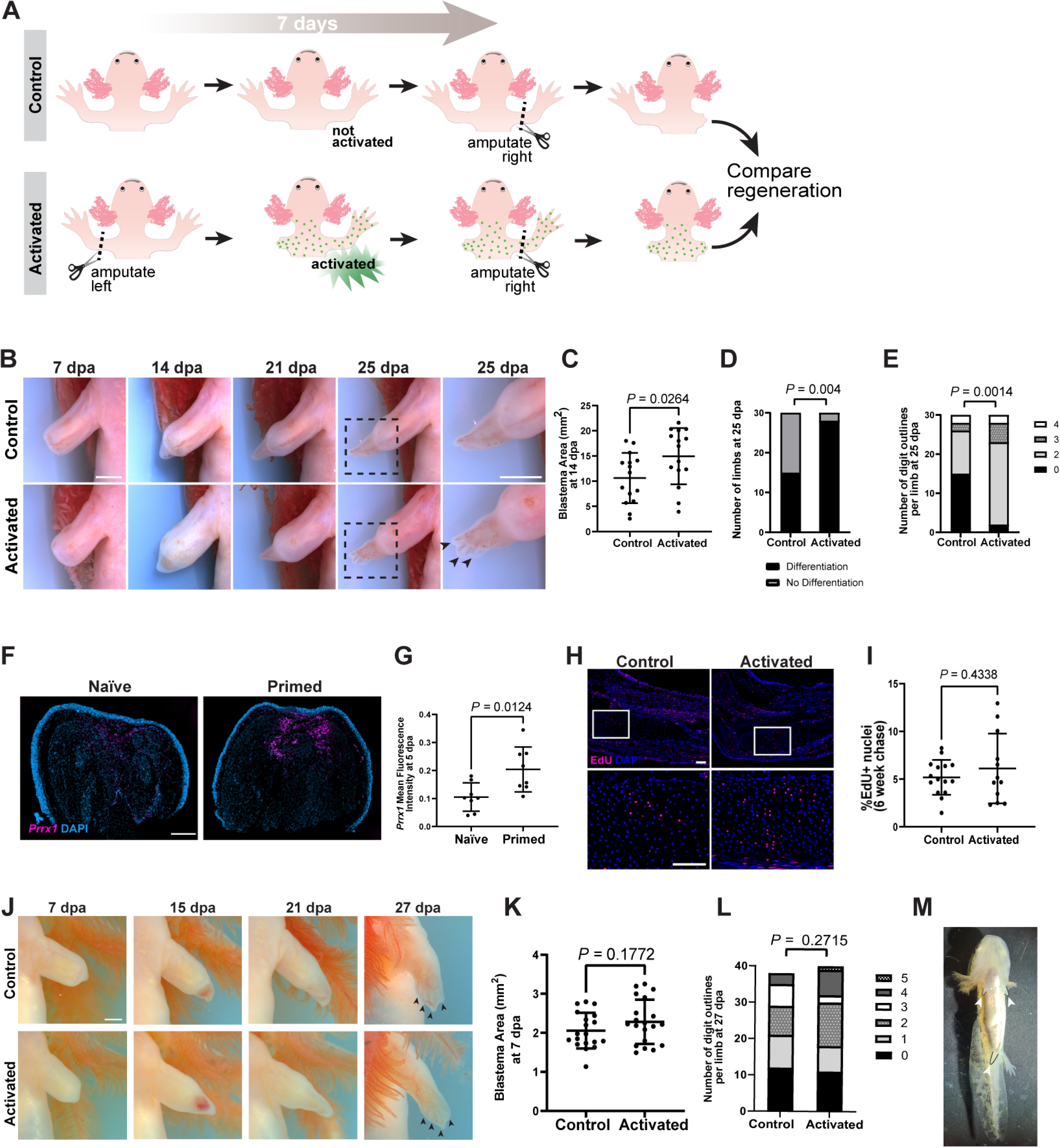
Systemically-activated cells are primed for future regeneration events. **(A)** Experimental design. We compared regeneration rates of systemically-activated and naïve, control limbs. Green dots, activated, EdU positive nuclei. **(B)** Representative images of regenerating systemically-activated or control limbs (ventral view). **(C-E)** Quantification of blastema area at 14 dpa **(C)**, number of differentiating limbs **(D),** and number of digit outlines **(E)** at 25 dpa (n=15 animals for each condition in blastema area measurement, n = 30 limbs for each condition in differentiation and digit outline quantification). **(F)** *Prrx1* HCR-FISH was performed on tissue sections at 5 dpa following local amputation of naïve limbs (no previous amputation) and primed limbs (previously amputated on the contralateral side 7 days earlier). **(G)** Quantification of *Prrx1* mean fluorescence intensity across entire tissue sections of naïve and primed blastemas (n = 8 animals for each group). **(H)** EdU and DAPI staining of intact control limbs (left) and limbs contralateral to amputation (right) harvested 6 weeks after EdU administration. **(I)** Quantification of percent EdU positive nuclei of intact and contralateral limbs in **(H),** (n = 15 animals for intact and n = 12 animals for contralateral group). **(J)** Representative images of systemically-activated and naïve limbs amputated 28 days post-contralateral amputation (ventral view, 7-21 dpa; dorsal view, 27 dpa). **(K-L)** Quantification of blastema area at 7 dpa **(K)** and number of digit outlines at 27 dpa **(L)** (n = 19 animals for control, n = 20 animals for activated for blastema area measurement, n = 38 limbs for control and n = 40 limbs for systemically-activated animals in digit outline quantification). **(M)** Tiger salamander housed in a near-naturalistic pond simultaneously regenerating three limbs (arrowheads). All data shown as mean ± SD. Statistical significance was determined using unpaired two-tailed Welch’s t-test in **(C)**, **(G)**, **(I)** and **(K)**; and Fisher’s exact test in **(D)**, and Fisher’s exact test with Freeman-Halton extension in **(E)** and **(L).** Scale bars, 2 mm in **(B)** and **(J)**; 500 µm in **(F)** and **(H).** dpa, days post-amputation. EdU, 5-ethynyl-2′-deoxyuridine.

We also investigated how many divisions cells undergo following their systemic activation and how long the priming effect lasts. Using a short-pulse EdU and long-chase strategy, we found most EdU-labeled clones in limbs to be composed of 2-8 cells (**Figure 1H**), indicating that amputation-induced proliferation may last only a few cell cycles. Concordantly, we found there to be no difference in the fraction of EdU+ cells by 6 weeks post-amputation (**Figure 1I**). Six weeks is enough time for larval axolotls to completely regenerate a limb. We also found that systemically-activated limbs challenged with local amputation 4 weeks after initial activation regenerated at the same rate as naïve limbs, indicating the effect stimulated by initial limb loss is relatively short-lived (**Figure 1J-L**), although subsequent distant injuries could further modify this timeframe.

We next considered whether there could be a need for wild salamanders to sequentially regenerate multiple limbs over a short timeframe. This is conceivable, as limb and tail regeneration following conspecific or predatory bite wounds is frequent in wild salamander populations^2,3^ and some salamander species even use a form of tail autotomy.^3^ Axolotls (*Ambystoma mexicanum*) are critically endangered in the wild^4^ and consequently could not be investigated in a natural setting. We therefore examined tiger salamanders (*Ambystoma tigrinum*), close cousins of axolotls, found in a naturalistic outdoor pond setting, for evidence of simultaneous multiple limb regeneration. We found that, indeed, many larval tiger salamanders had multiple limbs at different, off-set states of regeneration in this natural setting (**Figure 1M**; N>6), implying that the amputation injuries occurred sequentially rather than simultaneously.

Indeed, the loss of a first limb might be expected to reduce mobility and perhaps predispose an animal to subsequent predation. Additionally, loss of multiple limbs in individuals has been documented in wild salamander populations, such as eastern hellbenders (*Cryptobranchus alleganiensis alleganiensis*)^5^ and ocoee salamanders (*Desmognathus ocoee*).^6^ These data indicate that in certain settings, limb loss is likely to occur repetitively, and hence to be physiologically relevant for salamanders. Collectively, these data demonstrate that the initiation of limb regeneration can be positively influenced by previous limb loss elsewhere in the body, within a time frame relevant to when cells remain systemically activated and suggest that responses to repeated limb loss may be relevant in the wild.

### Sympathetic nervous system is required for systemic activation and priming for limb regeneration

Determining how amputation-induced signals spread throughout the body is important for unraveling the molecular and cellular mechanisms of the systemic activation response. We hypothesized that the peripheral nervous system (PNS) might be a conduit for this type of information, as it rapidly transduces sensory afferent information to the spinal cord and brain, as well as sends motor and autonomic efferent signals from the brain to the body, thereby facilitating communication between distant anatomical locations. We tested whether systemic activation requires peripheral innervation at either the responding site (**Figure 2A-E**) and/or the amputation site (**Figure 2F-G**) using surgical denervation via brachial plexus nerve transection or sciatic nerve transection and comparing it to sham operation controls (**Figure S1**). We found that systemic activation did not occur when these distant sites contralateral to amputation were denervated at the time of the amputation injury (**Figure 2A-B**). As a control, we found that systemic activation still occurred following amputation when these distant sites (contralateral limbs) underwent sham operations. We parsed the contralateral tissues into three broad tissue categories—epidermis, skeletal elements, and internal (non-skeletal, soft) tissues—and quantified the extent of systemic activation within each (**Figure 2C-E**). This analysis revealed that each of these three tissue types requires an intact nervous system for cellular activation. We also found that when limbs were denervated and then immediately amputated, the amputation no longer provoked systemic activation in contralateral limbs (distant sites) (**Figure 2F-G**). Sham surgeries in which limbs were not denervated prior to their amputation still provoked the systemic activation response. These results demonstrate that peripheral innervation of both the amputated limb and the distantly-responding limb is required for systemic activation of progenitor cells. Therefore, both neuronal input that senses injury and neuronal output that mediates efferent signaling to tissues could regulate systemic activation.

**Figure 2.**
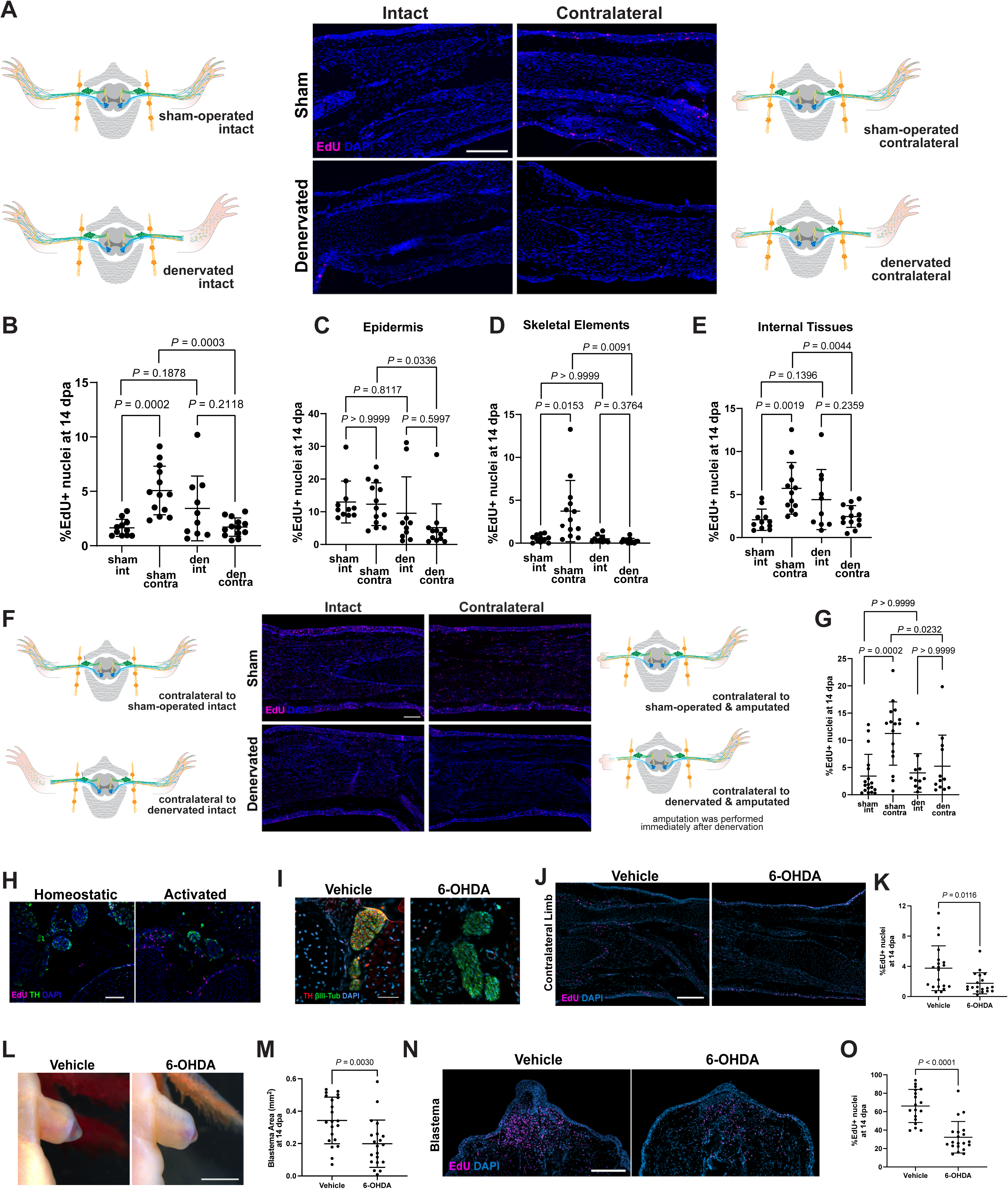
PNS at both amputation site and responding site are required for systemic activation. (**A-E**) Nerve bundles innervating limbs contralateral to unilateral amputations were transected or sham-operated, and animals were assayed for systemic activation. Animals receiving unilateral denervations or sham-operations with no amputations were used as controls. **(A)** EdU and DAPI staining of intact and contralateral limb tissue sections of denervated and sham-operated animals in the intact state or at 14 days post-amputation. **(B)** Quantification of percent EdU positive nuclei in denervated and sham-operated at 14 dpa. **(C-E)** Breakdown of tissue types represented in **(B)**. **(C)** Epidermis. **(D)** Skeletal elements. **(E)** Internal, non-skeletal tissues. (n = 11 animals for sham, n = 10 animals for denervated, n = 13 animals for sham amputated and denervated amputated), **(F-G)** Nerve bundles were transected or sham-operated in limbs that were immediately amputated thereafter. **(F)** EdU and DAPI staining of intact and contralateral limb tissue sections of denervated and sham-operated animals in the intact state or at 14 days post-amputation. **(G)** Quantification of percent EdU positive nuclei in denervated and sham-operated at 14 dpa (n = 18 animals for sham, n = 16 for sham amputated, n = 11 for denervated, and n = 12 for denervated amputated). **(H)** Representative image of cross-sectioned sympathetic nerve bundles (stained with α-TH) and EdU+ cells in adjacent tissues in homeostatic (left) and systemically-activated (right) samples. **(I)** Representative images of cross-sectioned nerve bundles of vehicle and 6-OHDA treated axolotl limbs. **(J)** EdU and DAPI staining of limbs contralateral to unilateral amputations treated with 6-OHDA or vehicle control, harvested at 14 days post-amputation. **(K)** Quantification of percent EdU positive nuclei in contralateral limbs in **(J)** (n = 21 animals for vehicle and n = 20 animals for 6-OHDA groups). **(L)** Representative regenerating limbs of 6-OHDA and vehicle-treated animals (ventral view). **(M)** Quantification of blastema area at 14 days post-amputation (n = 21 animals for vehicle and n = 20 animals for 6-OHDA groups). **(N)** EdU and DAPI staining of regenerating limbs treated with 6-OHDA or vehicle control, harvested at 14 days post-amputation. **(O)** Quantification of percent EdU positive nuclei in contralateral limbs in **(N)** (n = 21 animals for vehicle and n = 20 animals for 6-OHDA groups). All data shown as mean ± SD. Statistical significance was determined using unpaired two-tailed Welch’s t-tests with Bonferroni-Dunn correction in **(B)**, **(C)**, **(D)**, **(E)** and **(G)**, unpaired two-tailed Welch’s t-tests in **(K)**, **(M)** and **(O)**. Scale bars, 500 µm in **(A)**, **(F)**, **(J)** and **(N)**, 200 µm in **(H)**, 100 µm in **(I)**, and 2 mm in **(L)**. den, denervated. int, intact. contra, contralateral. 6-OHDA, 6-hydroxydopamine. EdU, 5-ethynyl-2′-deoxyuridine. See also **Figure S1**.

We next addressed which component of the peripheral nervous system regulates systemic activation. We hypothesized that limb loss is likely to provoke canonical stress pathways, such as activation of the sympathetic nervous system that mediates the “fight or flight” response. Supporting a possible role for sympathetic innervation, we noticed in our tissue sections that EdU+ cells were located near sympathetic nerves, identified by a tyrosine hydroxylase (TH) antibody (**Figure 2H**). We therefore tested whether sympathetic innervation is required for systemic activation. We administered 6-hydroxydopamine (6-OHDA) to axolotls to ablate sympathetic nerve processes and confirmed this effect in limb tissues using an antibody against TH (**Figure 2I**). Following 6-OHDA treatment, we performed limb amputation and assayed for systemic activation in contralateral limbs. We found that sympathetic nerve ablation blocks systemic activation (**Figure 2J-K**). Intriguingly, we also found that amputated limbs displayed significant impairment in blastema formation in the 6-OHDA treated animals (**Figure 2L-M**). Furthermore, these 6-OHDA treated limbs contained significantly fewer proliferating cells than vehicle controls (**Figure 2N-O**). These results demonstrate that sympathetic innervation is required for both systemic activation and for local limb regeneration. They also reveal that there may be mechanistic connections between systemic responses of limb cells to amputation and the ability of these cells to participate in localized regeneration. We therefore sought to further characterize the molecular effectors of noradrenergic signals in systemic activation and in limb regeneration.

### α_2A_ adrenergic receptor activity is required for systemic activation and priming

The primary adrenergic signaling effector of the sympathetic nervous system is noradrenaline, which acts as a neurotransmitter to provoke responses in peripheral tissues.^7^ Noradrenaline can bind to several different adrenergic receptors. Pharmacological inhibitors exist that antagonize specific noradrenaline-receptor interactions. We, thus, used these to address which adrenergic receptors are relevant to systemic activation and limb regeneration in axolotls. We first treated axolotls with yohimbine, a well-studied ADRA2A antagonist,^8,9^ to test for an adrenergic signaling requirement in both the regenerating limb and in systemic activation. We found that in animals treated with yohimbine, amputated limbs had lower numbers of proliferating cells (**Figure 3A-B**), and limbs contralateral to amputation did not undergo systemic activation (**Figure 3C-D**), demonstrating that adrenergic signaling is required for robust cell proliferation at the injury site and for systemic cell activation in response to amputation. We then asked whether adrenergic signaling is also required for priming distant tissues for future regeneration (**Figure 3E**). We found that when axolotls were treated with yohimbine from 2 days before to 7 days after amputation, priming was diminished (**Figure 3F**), with both initial blastema growth (**Figure 3G**) and limb differentiation (**Figure 3H**) occurring more slowly. We tested whether stimulating adrenergic signaling via the α_2A_-adrenergic agonist clonidine might enhance priming and found evidence that this pathway can indeed be stimulated to accelerate blastema growth of distant tissues that have been systemically activated (**Figure 3I-K**). Collectively, these data support a model in which noradrenaline released from sympathetic neurons provokes systemic activation through α_2A_-adrenergic receptors in target tissues, that this process is necessary for priming cells for faster future regeneration, and that targeting this pathway with a pharmacological agonist can enhance regenerative priming.

**Figure 3.**
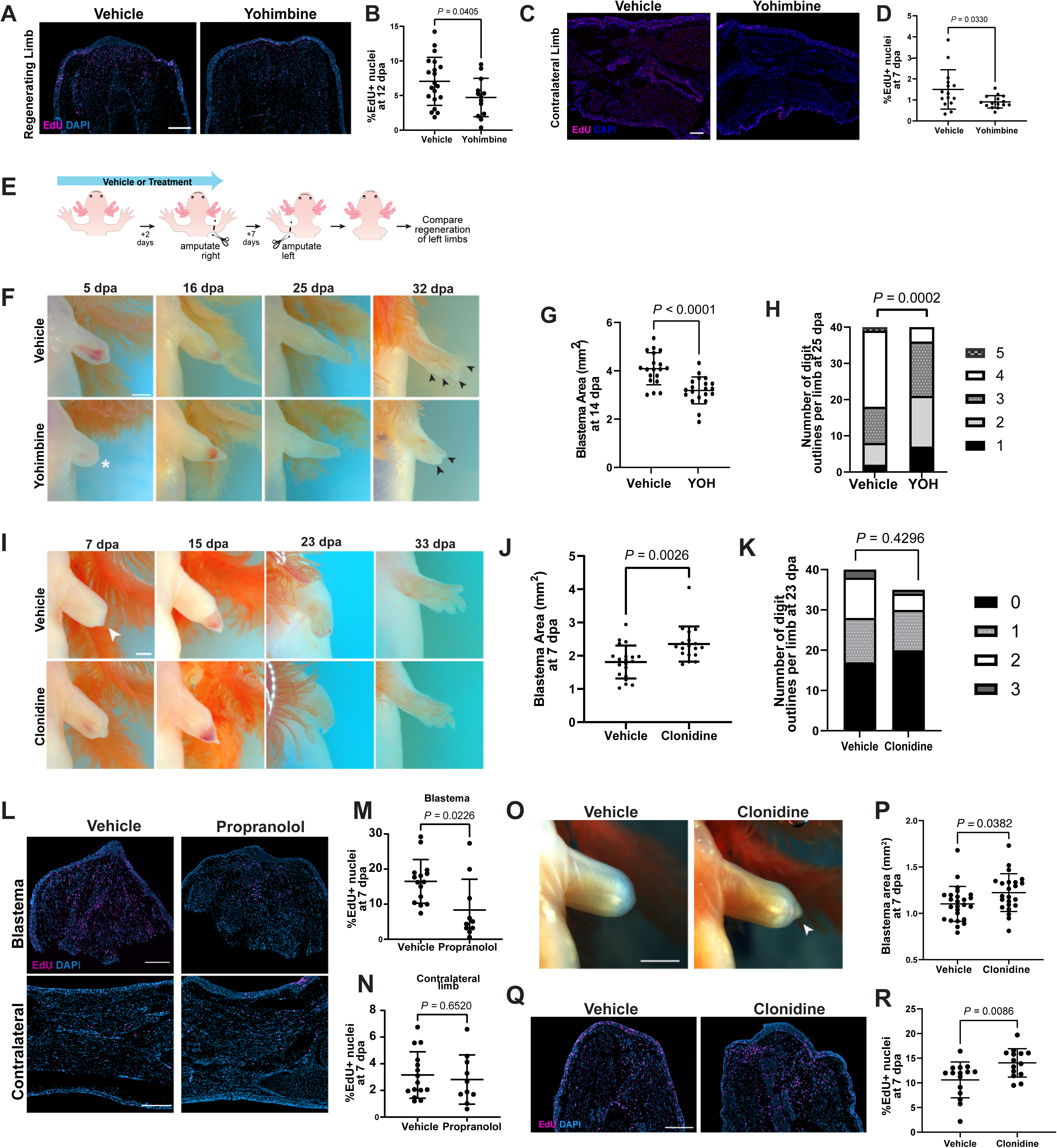
α-Adrenergic signaling is required for systemic activation and priming. **(A)** EdU and DAPI staining of regenerating blastemas from animals treated with yohimbine or vehicle control, harvested at 12 days post-amputation. **(B)** Quantification of percent EdU positive nuclei in blastemas in **(A)** (n = 20 animals for vehicle, n = 13 animals for yohimbine). **(C)** EdU and DAPI staining of limbs contralateral to unilateral amputations treated with yohimbine or vehicle control, harvested at 7 days post-amputation. **(D)** Quantification of percent EdU positive nuclei in contralateral limbs in **(D)** (n = 15 animals for each group). **(E)** Experimental design. Animals were treated with chemical agonists or antagonists (treatment) or vehicle solution by daily injections or water treatments starting from 2-3 days before to 7 days post unilateral amputations. The treatments were then ceased, and the uninjured contralateral limbs were unilaterally amputated. The regeneration rates of the second set of amputations were compared between treatment groups. **(F)** Representative images of regenerating contralateral limbs of yohimbine and vehicle solution-treated animals (ventral view, 5-25 dpa; dorsal view, 32 dpa). **(G)** Quantification of blastema area at 14 days post-amputation (n=15 animals for each condition). **(H)** Quantification of the number of digit outlines at 25 days post-amputation (n = 30 limbs for each condition). **(I)** Representative images of regenerating contralateral limbs of clonidine and vehicle solution-treated animals (ventral view, 7-23 dpa; dorsal view, 33 dpa). **(J)** Quantification of blastema area at 7 days post-amputation. (n = 20 animals for vehicle and n = 18 animals for clonidine). **(K)** Quantification of the number of digit outlines at 23 days post-amputation (n = 40 limbs for vehicle and n = 35 limbs for clonidine). The same group of vehicle control animals was used in Figure 4 (I-K). **(L-N)** Animals were treated with beta-adrenergic receptor antagonist propranolol starting from 2 days before to 7 days post unilateral amputations. **(L)** EdU and DAPI staining of blastemas and limbs contralateral to unilateral amputations treated with propranolol or vehicle control, harvested at 7 days post-amputation. **(M)** Quantification of percent EdU positive nuclei within the blastema area of the initially-amputated limbs (n = 15 animals for vehicle and n = 10 for propranolol). **(N)** Quantification of percent EdU positive nuclei within contralateral limbs at 7 dpa (n = 15 animals for vehicle and n = 10 for propranolol). **(O-R)** Animals were treated with Adra2a agonist clonidine or vehicle solution starting from 2 days before to 7 days post unilateral amputations. **(O)** Representative images of regenerating limbs at 7 days post-amputation. **(P)** Quantification of blastema area at 7 dpa (n = 25 animals for vehicle and n = 20 animals for clonidine). **(Q)** Representative images of EdU staining of regenerating limbs harvested at 7 days post-amputation. **(R)** Quantification of percent EdU positive nuclei within the blastema area (n = 15 animals for vehicle and n = 14 for clonidine). All data shown as mean ± SD. Statistical significance was determined unpaired two-tailed Welch’s t-tests in **(B)**, **(D)**, **(G)**, **(M)**, **(N), (P)** and **(R)**; Welch’s ANOVA with Dunnett’s T3 correction in **(J)**; Fisher’s exact test with Freeman-Halton extension in **(H)** and **(K)**. Scale bars, 500 µm in **(A), (C), (L)** and **(Q)**; 2 mm in **(F), (I)** and **(O)**. hpa, hours post-amputation. dpa, days post-amputation. EdU, 5-ethynyl-2′-deoxyuridine. YOH, yohimbine. See also **Figure S2.**

We also assessed possible contributions to systemic activation from other adrenergic receptors that can also bind noradrenaline. We used an identical treatment regimen to administer the non-selective β-adrenergic receptor antagonist propranolol^10^ to axolotls (**Figure 3L-O**). We found that while propranolol treatment led to a significantly decreased fraction of cells in S-phase within blastemas of regenerating limbs (**Figure 3L-M**), it had no effect on systemic activation (**Figure 3L, N**). These results indicate β-adrenergic signaling is only required for the localized cellular proliferation response at the amputation site.

We next asked whether stimulating α_2A_-adrenergic signaling in the context of a single limb amputation alone is sufficient to promote faster blastema formation, and, using clonidine treatment, we found that this was the case (**Figure 3O-P**). This enhanced regeneration response is accompanied by an increased fraction of proliferating cells in the amputated limbs (**Figure 3Q-R**). We also tested whether clonidine administration is sufficient to promote systemic activation in the absence of amputation injury and found it to be insufficient (**Figure S2B-C**). Clonidine treatment also could not rescue systemic activation when either amputated limbs (**Figure S2D-E**) or responding limbs (**Figure S2F-G**) were denervated. Together, these results argue that systemic activation and priming require α-adrenergic signaling, and that stimulating this pathway is sufficient to promote faster limb regeneration, albeit likely synergizes with other pathways downstream of nerves. These findings also argue that β-adrenergic signaling is required specifically at the injury site to promote limb regeneration.

### mTOR signaling is required for systemic activation in axolotl

Our findings highlight the sympathetic nervous system and adrenergic signaling as key drivers of systemic priming and proliferation after amputation in axolotls. A similar systemic response occurs in mice after local injury whereby distant cells re-enter the cell cycle.^11^ In mice, this activation is linked to circulating HGFA (hepatocyte growth factor activator), which activates HGF (hepatocyte growth factor) and stimulates mTOR signaling.^11,12^ Additionally, a recent study of the amputation site identified a key role for mTOR in translational regulation of the wound-healing stage of axolotl limb regeneration.^13^ We investigated the potential participation of mTOR in systemic activation in axolotl. We found that axolotls treated with the mTOR inhibitor rapamycin^14^ regenerate limbs more slowly (**Figure 4A-C**), with lower numbers of proliferating cells in the blastema (**Figure 4D-E**), and that both systemic activation (**Figure 4F-G**) and priming are inhibited by rapamycin treatment (**Figure 4H-K)**. Considering the role of mTOR in nutrient sensation, cell growth, and proliferation,^15^ we tested whether fasted axolotls exhibit altered levels of systemic activation and whether cell proliferation in this context might additionally be regulated by mTOR signaling. We found that global cell proliferation rates in amputated and fasted axolotls were dramatically lower than typically observed in well-fed animals (**Figure 4L-M**). Yet, we also found that in the context of an amputation, the number of proliferating cells at distant sites still reduced by inhibition of mTOR signaling (**Figure 4L-M).** These results demonstrate that mTOR signaling is important for systemic activation responses to limb amputation in axolotl.

**Figure 4.**
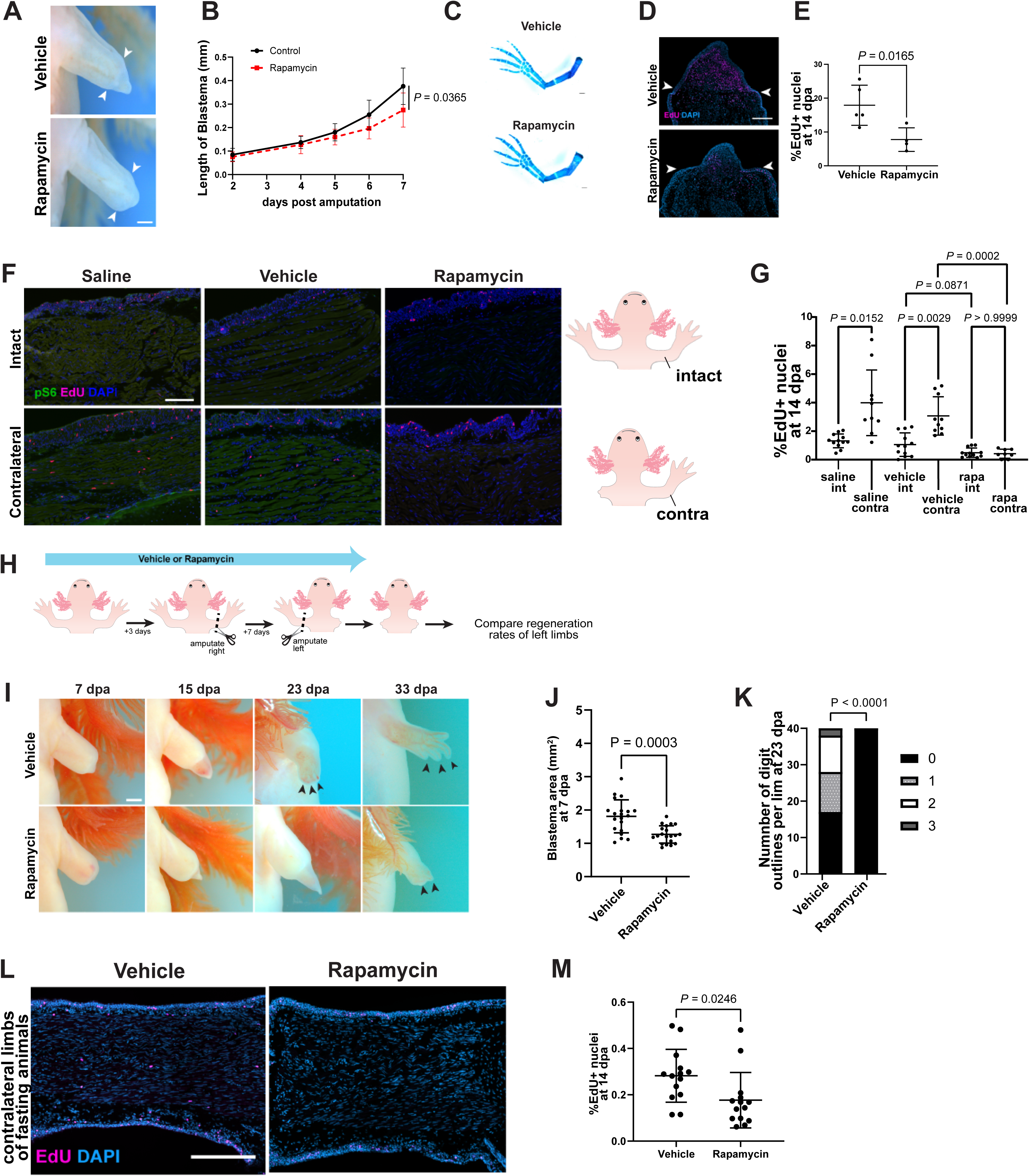
mTOR signaling is required for systemic activation and priming. **(A)** Regenerating limbs at 7 dpa treated with rapamycin or vehicle control. Arrowheads mark the amputation plane. **(B)** Blastema growth curves of rapamycin or vehicle-treated animals (n = 10 animals per group). Statistical significance was determined using unpaired two-tailed Welch’s t-test with Holm-Šídák correction). **(C)** Representative skeletal preparations of regenerated limbs with Alcian blue and Alizarin red at 10 weeks post-amputation. **(D)** EdU and DAPI staining of rapamycin or vehicle-treated blastemas at 14 dpa. Arrowheads mark the amputation plane. **(E)** Quantification of percent EdU positive nuclei in **(D)**. **(F)** pS6 and EdU staining of intact (homeostatic) and contralateral limb tissue sections from animals treated with saline, vehicle, or rapamycin at 14 dpa. **(G)** Quantification EdU positive nuclei at 14 dpa (n = 12 limbs for intact saline, intact vehicle and intact rapamycin groups, n = 10 limbs for amputated saline and amputated vehicle, and n = 8 limbs for amputated rapamycin group). **(H)** Experimental design to test requirement for mTOR in priming. Animals were treated with rapamycin or vehicle solution by daily injections starting from 3 days before to 7 days post unilateral amputations. The treatments were then ceased, and the uninjured contralateral limbs were unilaterally amputated. The regeneration rates of the second set of amputations were compared between treatment groups. **(I)** Representative images of regenerating contralateral limbs of rapamycin and vehicle solution-treated animals. **(J)** Quantification of blastema area at 7 days post-amputation. **(K)** Quantification of the number of digit outlines at 23 days post-amputation (n=20 animals for vehicle and rapamycin; n = 40 limbs for vehicle and rapamycin for each condition in digit outline quantification). The same group of vehicle control animals were used in Figure 3 (I-K). **(L-M)** Animals fasting for four weeks were treated with mTORC1 inhibitor rapamycin or vehicle control solution from 2 days before to 14 days post-unilateral amputations. **(L)** EdU and DAPI staining of limbs contralateral to unilateral amputations from animals treated with rapamycin or vehicle control, harvested at 14 dpa. **(M)** Quantification of percent EdU positive nuclei in **(L)**. All data shown as mean ± SD. Statistical significance was determined using unpaired two-tailed Welch’s t-test with Holm-Šídák correction in **(B)**; unpaired two-tailed Welch’s t-test in **(E)** and **(M)**; unpaired two-tailed Welch’s t-test with Bonferroni-Dunn correction in **(G)**; two-tailed Welch’s ANOVA with Dunnett’s T3 correction for multiple hypothesis testing in **(J)**; and Fisher’s exact test with Freeman-Halton extension in **(K)**. Scale bars, 0.5 mm in **(A)**; 500 µm in **(D)**, **(F)** and **(L)**; 2 mm in **(I)**. rapa, rapamycin. Int, intact. contra, contralateral. dpa, days post-amputation. pS6, phospho-S6 ribosomal protein; EdU, 5-ethynyl-2′-deoxyuridine.

### Adrenergic signaling operates upstream of mTOR signaling in systemic activation and limb regeneration

Since we found that mTOR signaling is required for systemic activation in axolotls, we sought to determine whether this dependency is mechanistically related to the requirement for adrenergic signaling in systemic activation and limb regeneration. We inhibited α_2A_-adrenergic signaling using yohimbine, amputated limbs, and assayed for mTOR signaling in contralateral limbs using pS6 as a readout. We found that yohimbine treatment significantly reduced mTOR signaling in contralateral limbs, providing evidence that mTOR is under the control of α_2A_-adrenergic signaling in the systemic response to amputation (**Figure 5A-B**). In separate experiments, we treated naïve axolotls with the yohimbine or with the β-adrenergic antagonist propranolol, amputated limbs, and assayed for mTOR signaling in each case using pS6 as a readout. We found that both yohimbine treatment (**Figure 5C-D)** and propranolol treatment (**Figure 5E-F**) significantly reduced mTOR signaling in stump tissues. These data argue that adrenergic signaling is required for mTOR signaling in the context of axolotl limb regeneration. To further investigate mTOR signaling dependency on adrenergic signaling, we leveraged an existing axolotl limb fibroblast cell line, AL-1 cells.^16^ We treated AL-1 cells with adrenaline and found it was sufficient to significantly enhance mTOR signaling *in vitro* (**Figure 5G-H**). Collectively, these data argue that axolotl limb cells use adrenergic signals to upregulate mTOR signaling.

**Figure 5.**
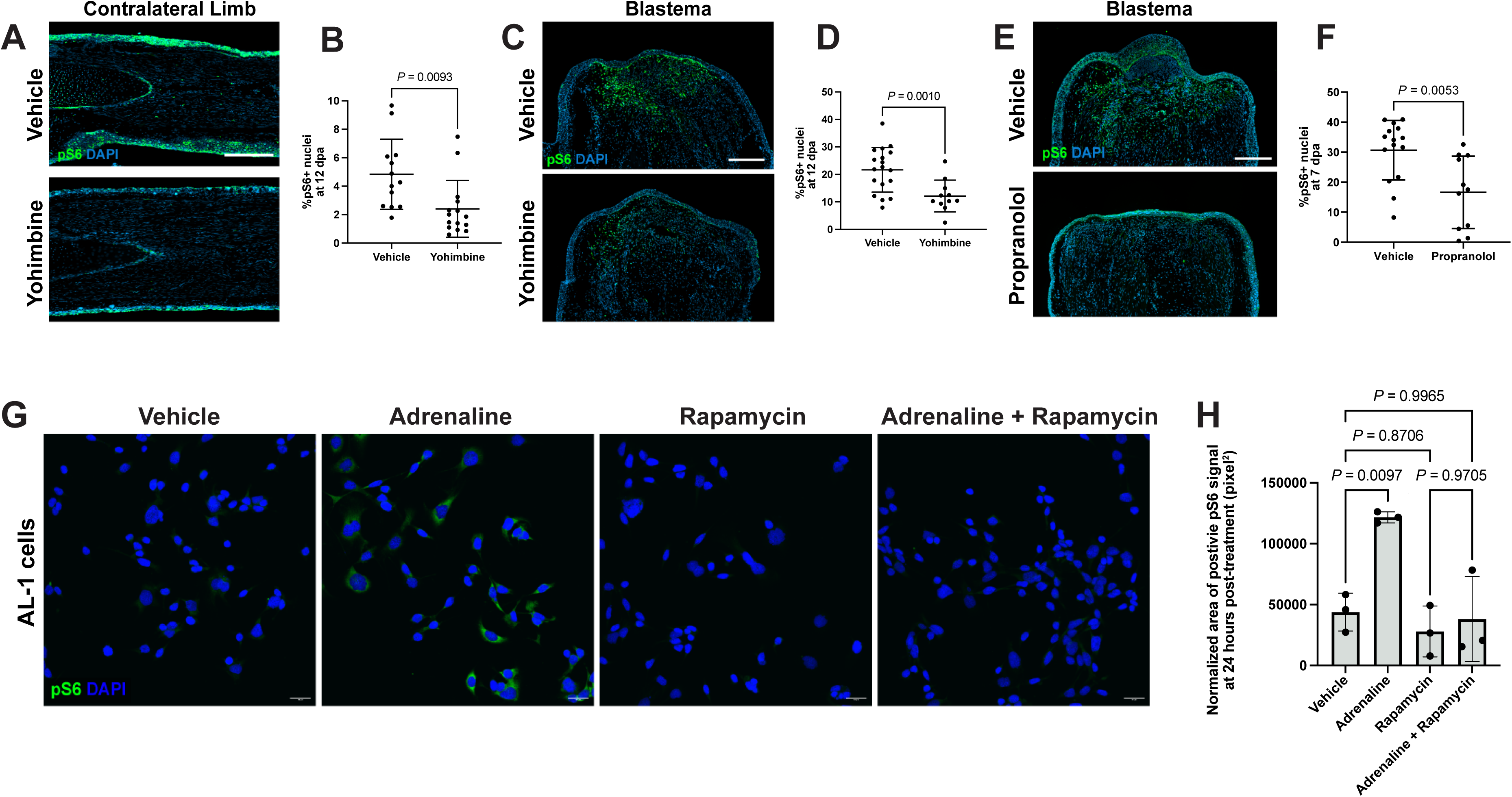
Adrenergic signaling operates upstream of mTOR signaling in systemic activation and limb regeneration. **(A)** pS6 and DAPI staining of limbs contralateral to amputation from animals treated with yohimbine or vehicle control, harvested at 12 days post-amputation. **(B)** Quantification of percent pS6 positive cells in limbs in **(A)** (n = 13 animals for vehicle, n = 15 animals for yohimbine). **(C)** pS6 and DAPI staining of regenerating blastemas from animals treated with yohimbine or vehicle control, harvested at 12 days post-amputation. **(D)** Quantification of percent pS6 positive cells in blastemas in **(C)** (n = 18 animals for vehicle, n = 11 animals for yohimbine). **(E)** pS6 and DAPI staining of regenerating blastemas from animals treated with propranolol or vehicle control, harvested at 7 days post-amputation. **(F)** Quantification of percent pS6 positive cells in blastemas in **(E)** (n = 15 animals for vehicle, n = 11 animals for propranolol). **(G)** pS6 and DAPI staining of AL-1 cells treated with adrenaline, rapamycin or vehicle solution for 24 hours. **(H)** Normalized area of positive pS6 signal of AL-1 cells shown in **(G)**. Data shown as mean ± SD. Statistical significance was determined using unpaired two-tailed Welch’s t-tests in **(B)**, **(D)** and **(F)**; ordinary one-way ANOVA with Dunn-Šidák correction for multiple hypothesis testing in **(H)**. Scale bars, 500 μm in **(A)**, **(C)** and **(E)**; 50 µm in **(G)**. pS6, phospho-S6 ribosomal protein. dpa, days post-amputation.

### Systemically-activated cell types are shared with proliferating cell types in homeostasis and are epigenetically distinct

We sought to identify the types of cells that become systemically activated and the signals they might use to communicate with other cells in activated limbs. In our earlier study, we identified a subset of systemically-activated cells (SACs) as satellite cells^1^, which serve as stem cells for muscle regeneration in many species, including axolotls.^17^ However, the identities of most axolotl SACs remained unknown. Moreover, commercially-available antibodies recognizing axolotl marker proteins show limited success. Here, to overcome this limitation, we designed a marker-agnostic FACS strategy to purify and profile the transcriptomes of individual SACs based on their 4C nuclear content compared to 2C content of non-proliferative cells (**Figure 6A-B**). We profiled transcriptomes from 4C SACs and 2C cells from both systemically-activated and naïve control limbs. We merged these data and classified cell types across samples (**Figure 6C, Figure S3, Table S1, Table S2**). The vast majority of proliferative 4C cell types had correlating clusters among non-proliferative 2C cells (**Figure 6D**). Furthermore, the SAC types overlap with 4C cell types collected from intact, naïve limbs (**Figure 6E**). These data indicate that SACs largely represent stem and progenitor cell types that ordinarily cycle during homeostasis, including during organismal growth and wear-and-tear repair. This finding suggests that regenerative strategies could be developed to potentially steer the activities of pre-existing progenitor cell types from growth and homeostatic functions toward building a blastema including, conceivably, in mammals.

**Figure 6.**
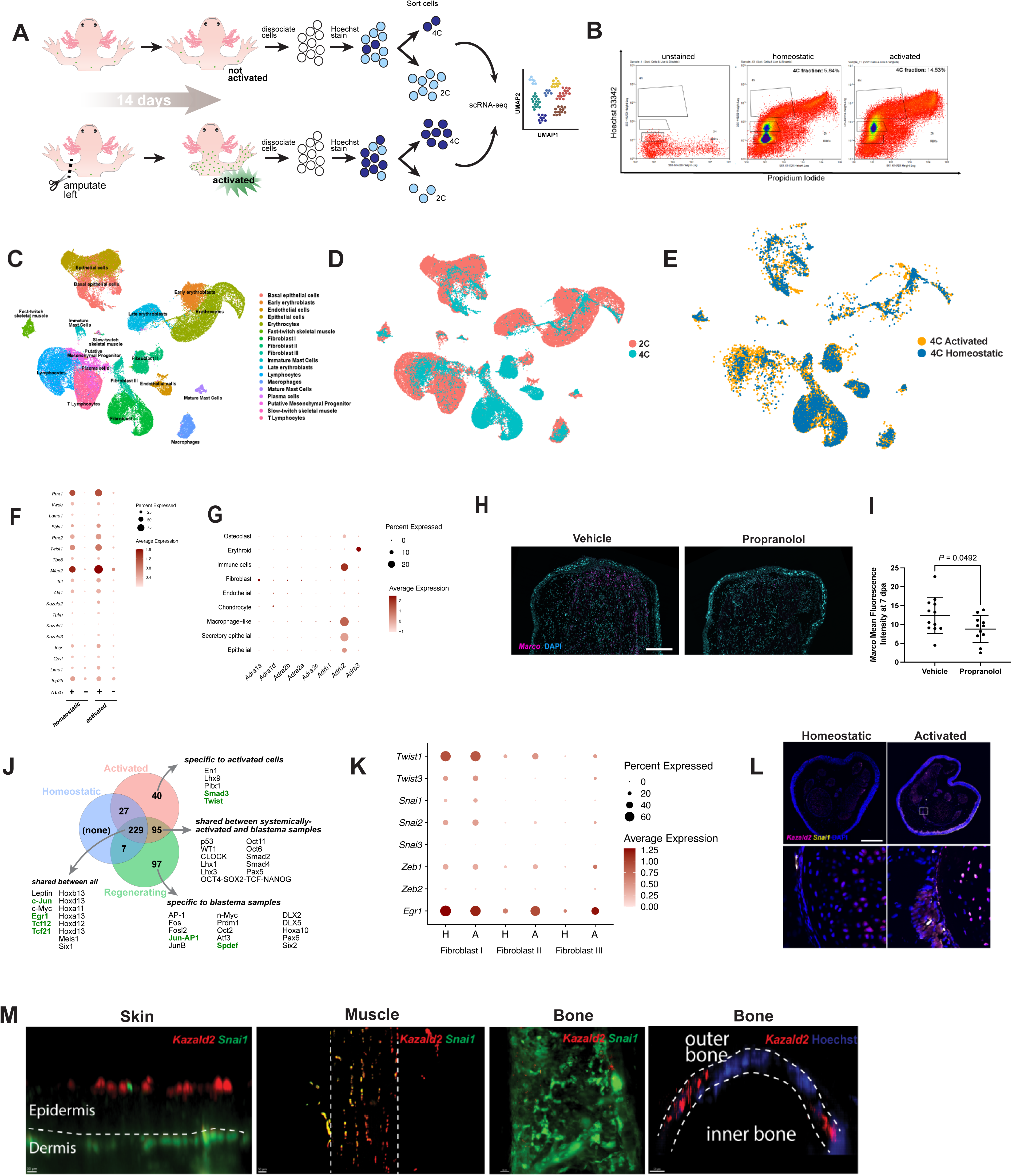
Systemically-activated cells represent the same broad cell types as cells proliferating during homeostasis and are epigenetically primed. **(A)** Experimental design to identify systemically activated cells and compare to proliferating cells during homeostasis. Dividing (4C) and non-dividing (2C) cells of systemically-activated and naïve, control limbs were isolated by sorting dissociated cells according to their DNA content. Each fraction was sequenced separately by single cell RNA-sequencing. **(B)** Representative FACS distribution following Hoechst stain of homeostatic and systemically-activated limb cell samples. **(C)** UMAP plot of cells collected from homeostatic control limbs (n = 4) and systemically-activated limbs (n = 4). **(D)** UMAP plot of cells harvested from homeostatic and activated limbs, with 2C cells colored in red and 4C cells colored in blue **(E)** UMAP plot of 4C cells with cells harvested from the homeostatic limb in blue and systemically-activated limb in yellow. **(F)** Dot plot showing the relative frequency of expression (dot size) and expression level (color) for differentially expressed blastema-specific genes across *Adra2a*+ and *Adra2a*-cells in homeostatic (H) versus activated (A) samples. **(G)** Dot plot showing the relative frequency of expression (dot size) and expression level (color) for alpha– and beta-adrenergic receptors in regenerating axolotl limbs (7 timepoints from 3 hpa to 33 dpa combined), sequencing data re-analyzed from Li *et al.*^46^ (H) *Marco* HCR-FISH on blastema sections of propranolol or vehicle-treated animals **(I)** Quantification of *Marco* mean fluorescence intensity across blastema tissue sections of propranolol or vehicle-treated animals at 7 dpa (n = 12 animals vehicle, n = 11 for propranolol groups). **(J)** Venn diagram illustrating significantly enriched predicted transcription factor binding motifs within open chromatin of each of the three conditions sampled, 4C cells from homeostatic limbs, 4C cells from regenerating limbs (blastema cells), 4C systemically-activated cells. Representative transcription factors are noted, factors directly associated with EMT in green. **(K)** Dot plot showing the relative frequency of expression (dot size) and expression level (color) for epithelial-mesenchymal transition markers in homeostatic and activated fibroblast clusters. (L) *Kazald2 and Snai1* HCR-FISH on homeostatic and activated limb tissue sections. (M) *Kazald2* and *Snai1* RNAscope-FISH on wholemount homeostatic limbs. All data shown as mean ± SD. Statistical significance was determined using unpaired two-tailed Welch’s t-test in **(I).** Scale bars, 500 µm in **(H)** and **(L).** dpa, days post-amputation. See also **Figure S3, S4** and **S5, Table S1, S2** and **S3** and **Supplementary Movie 1.**

We also used these transcriptome data to identify cell types that express adrenergic receptors. We found that in both homeostatic (from uninjured animals) and activated limbs, fibroblasts are the primary cell type expressing *Adra2a* transcripts (**Figure S4A**), while β-adrenoreceptor transcripts are expressed by lymphocytes, mast cells, macrophages and endothelial cells in both homeostatic and activated limbs (**Figure S4B**). This data supports a model in which limb fibroblasts receive norepinephrine signals, potentially act as sensors of distant amputation. We then wondered if cells that express *Adra2a* might be primed toward a more regenerative state compared to other limb cells. We therefore computationally isolated *Adra2a*^+^ cells from our dataset and performed co-expression analyses with established blastema-enriched genes. We found that *Adra2a*^+^ cells in both homeostatic and activated limbs are enriched for expression of a suite of such genes, including *Prrx1*, *Prrx2*, *Vwde*, *Kazald2*, and *Twist1*, among others (**Figure 6F**). These data indicate that *Adra2a*-expressing cells are likely primed toward blastema cell states even before distant injury and that distant amputation provokes upregulation of genes that typically rise during *bona fide* blastema creation on amputated limb stumps.

We next sought to relate our findings about adrenoreceptor expression to cellular events regulating localized blastema formation within amputated limb stumps. We interrogated a published single-cell RNA-sequencing (scRNA-seq) dataset^18^ that profiled cells from homeostatic limbs and from limb stumps at several time points post-amputation. Intriguingly, we found the most significantly expressed adrenoreceptor to be *Adrb2*, which was expressed in several types of immune cells, including macrophages, as well as in epithelial cell populations of wound epidermis (**Figure 6G**). These results reinforce our earlier experimental finding that β-adrenergic signaling is important for local blastema formation and limb regeneration even though it is dispensable for systemic activation. Macrophages are known to be required at the amputation site for successful blastema formation and limb regeneration^19^, but the upstream signals that recruit them to the injury site are largely unknown. We hypothesized that adrenergic signaling and specifically epinephrine-mediated signaling through β-adrenoreceptors may be required for macrophage recruitment at the amputation site. We tested this hypothesis by blocking β-adrenergic signaling through propranolol administration and assayed for macrophages at the amputation site using HCR RNA *in situ* hybridization for the macrophage marker *Marco* (**Figure 6H**). We discovered that propranolol-treated samples harbored significantly fewer *Marco*^+^ macrophages at the amputation site (**Figure 6I**). A previous report demonstrated that immune cytokine IL-8 can promote recruitment of myeloid cells to regenerating axolotl limbs^20^, and in human monocytes, β-adrenergic signaling in myeloid cells can increase IL-8 release.^21^ These results support a model whereby β-adrenergic signaling promotes limb regeneration by controlling macrophage recruitment and/or proliferation during the blastema-forming stages of regeneration.

While the above described differences in gene expression could be important, we also recognized that epigenetic differences may contribute to systemic activation and the behaviors of SACs. To explore the chromatin accessibility profiles of SACs, we performed bulk ATAC-seq on SACs and 4C cells from homeostatic tissue samples as well as blastema tissues, permitting us to compare proliferating cells across all three contexts. Intriguingly, we uncovered 95 transcription factor binding motifs enriched in regions of open chromatin in both SACs and blastema cells but not in 4C cells from homeostatic, naïve axolotls **(Figure 6J, Table S3)**. Among the open chromatin regions in 4C cells, we found several that harbor predicted binding motifs for transcription factors that have direct roles in regulating epithelial-mesenchymal transition (EMT) in other contexts, such as *Twist*, *Egr1*, *Etv1*, *Tcf12*, *Tcf21*, *Spdef*, *Smad3*, *Jun* and *AP-1*. EMT-like processes are involved in wound healing, blastema formation, cell migration and tissue re-patterning during limb regeneration.^22,23^ Additionally, in other contexts, mTOR signaling interacts with EMT processes.^24^ We therefore reasoned that epigenetic regulation of these pathways in SACs could be important in priming them for future local regeneration. To investigate this possibility, we leveraged our scRNA-seq of homeostatic and systemically-activated limbs to explore the activity of EMT pathways. We particularly focused on fibroblasts, as they are an important source of many limb blastema cells.^25–28^ We found that several EMT-related transcription factors were upregulated in fibroblasts of the systemically-activated limb compared to the homeostatic limb **(Figure 6K).** Among these were *Egr1*, *Twist1*, *Snai1*, *Snai2* and *Zeb1*. In our transcription factor binding site analysis, we also found that Egr1 binding sites were shared across homeostatic, activated and regenerating limbs, whereas Twist binding sites were unique to activated limbs **(Figure 6J)**.

To visualize the expression of EMT regulators with respect to proliferating cells during homeostasis and systemic activation, we detected expression of *Snai1*. In our dataset, *Snai1* is upregulated in the Fibroblast I cluster upon systemic activation **(Figure 6K)**. We co-localized *Snai1* signal with *Kazald2*, a unique marker of this fibroblast cluster, and known for its regulation of limb regeneration.^29^ We found that in both homeostatic and activated samples (**Figure 6L**), some cells localizing to cartilage, perichondrium, nerve bundles, and muscle express *Kazald2*, *Snai1*, or both. In homeostatic samples, we observed the strongest co-expression signal for *Kazald2* and *Snai1* in the peri-skeletal tissue region (**Figure 6L**). Upon distant amputation injury, we observed a marked increase in expression of both *Kazald2* and *Snai1* (**Figure 6L**). We further looked at localization of *Kazald2* and *Snai1* transcripts in a wholemount limb and confirmed strong co-expression of the two genes in the epidermis and peri-skeletal tissues (**Figure 6M, Figure S5** and **Supplementary Movie 1)**. The findings suggest that EMT-like processes may be initiated at the epigenetic and transcriptional level by systemic activation. Interestingly, when we performed a short EdU pulse followed by a long chase (**Figure 1H**), we found that the resulting 2–8 cell clones in the systemically-activated limb remained clustered within their niches, suggesting that additional mechanisms restrict EMT-like progression until a local injury cue is present.

Collectively, our data demonstrate global differences in chromatin accessibility between homeostatic and systemically-activated states. They further provide evidence that SACs in uninjured tissues bear molecular resemblance to *bona fide* blastema cells, which may form a cellular basis for faster response in systemically-activated animals to future local injuries.

## DISCUSSION

Our results highlight a previously-unappreciated role for body-wide activation of tissue-resident cell types that cycle during organismal homeostasis and their repurposing for pro-regenerative responses during vertebrate limb regeneration. Our work indicates that the initiation phase of limb regeneration engages much more of the animal than previously appreciated and consists of multiple steps that are separable in both space and time. Nearly all previous work investigating regeneration initiation has focused exclusively on effects at the local injury site. However, emerging new data shows that during axolotl tail regeneration, immediate early response genes are activated in the brain.^30^ We previously demonstrated that axolotls respond to limb amputation by systemically activating a subset of cells in distant tissues to re-enter the cell cycle but did not address the mechanism of the response or its role in limb regeneration. The present study demonstrates that systemic activation is a priming step in the regenerative process on which subsequent cell behaviors necessary for blastema creation and, ultimately, regeneration are built. These results indicate an EMT-like event is likely the priming mechanism in distant tissues. This EMT event could tip cells toward regeneration by enabling their faster mobilization out of tissue niches after future local injury. These findings indicate that connecting global events to local signaling and cell behaviors is crucial to understanding the regeneration process.

We have here established that systemic stem cell activation is a key feature of the early regenerative response, highlighting the importance of understanding how this process is stimulated by amputation. These data support a model in which the mechanisms of systemic activation require peripheral innervation at both the injury and distant responding sites (**Figure 7A**) and define a primary role for sympathetic innervation in the initial stimulation of progenitor cells to re-enter the cell cycle. This role for the PNS is complementary to the previously-appreciated roles for the PNS as a source of mitogens in growing nascent blastemas larger^31–35^ and as a positive regulator of limb size during regeneration.^36^ At a mechanistic level, we uncover adrenergic signaling as a critical mediator of both systemic activation and priming for future regeneration.

**Figure 7.**
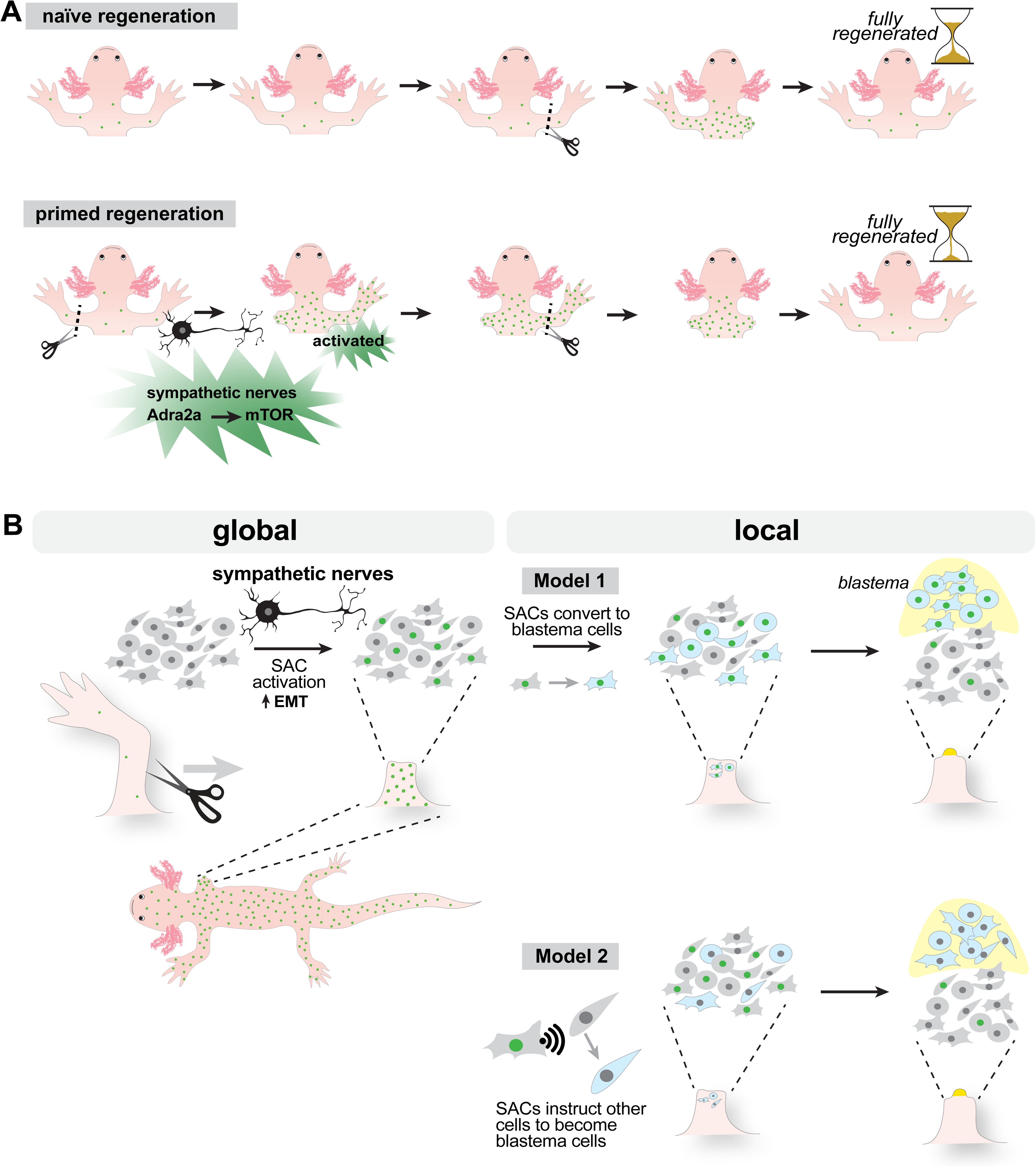
Main findings and implications of the study. **(A)** Systemic stem cell activation following amputation promotes faster regeneration. Systemic activation is driven by nerves and requires adrenergic signaling. **(B)** Two possible models for how systemic activation may promote local limb regeneration. In both models, some limb cells (green nuclei) re-enter the cell cycle (activation) and begin to proliferate in response to amputation. This step is systemic, occurring throughout the body, and common to both models. In the first model (top), some subset of activated progenitor cells later converts to the blastema cell state (blue cells). We postulate this step is local to the site of amputation. The subsequent migration of blastema cells to the distal-most tip of the stump creates a visible blastema (yellow). In the second model (bottom), systemically-activated cells signal to nearby cells to promote functions necessary for blastema formation, which could include dedifferentiation and migration. These models are not mutually exclusive.

We show that cells distant to amputation undergo proliferation while changing chromatin accessibility and gene expression, but there are several models that may explain the regenerative role of systemically-activated cells after activation. In one model (**Figure 7B**, model 1), they may go on to become blastema cells, acting as progenitor cells for new tissues. *In vivo* lineage tracing will be necessary to fully evaluate that possibility. In another model (**Figure 7B**, model 2), systemically-activated cells promote regeneration by acting as signaling hubs that influence the activity of other cells, perhaps priming them for regeneration. These models are not mutually exclusive, and possibilities for systemically-activated cells arising from traditional stem (or committed progenitor) cells or from dedifferentiation-type processes must be further investigated.

Our findings may provide potential insights into the evolution of regeneration by illustrating the possible advantage of systemic activation being a primary response to amputation. While this initial global activation event poises cells at the injury site toward a regenerative response, it simultaneously poises other tissues in the body toward accelerated regeneration should it become necessary. This global priming could have direct relevance in the wild for animals faced with intense predation—either from heterospecific or conspecific predation. Indeed, this might already have been of relevance in the stem lineage of modern amphibians some 290 Ma ago, as fossil evidence not only suggests that these ancient amphibians were capable of regeneration, but also that regeneration had occurred in more than one limb in single specimens.^2^

Axolotls are neotenic, and, in the wild, they live their entire lives in the water; however, this neotenic life cycle is a derived trait.^37^ For most species of salamanders, eggs are laid at very high density in a short time frame (for example, in vernal pools in the spring), resulting in high densities of larval salamanders and cannibalism.^38^ Because of these circumstances, and in conjunction with our direct evidence, salamanders may need to regenerate multiple limbs in a short time frame, especially if these limbs are necessary for more mature salamanders to leave the water for their terrestrial habitats. Thus, though outside the scope of this study, evaluating the potential adaptive value of systemic activation to salamanders in a natural setting, while a tantalizing possibility, will require controlled experimentation, including fieldwork. The possibility that systemic activation initiates limb regeneration in wild salamanders while also preparing other appendages for future regeneration is salient.

Several other outstanding questions remain for understanding the implications of our findings. We postulate that while the initial proliferative response to amputation is global in axolotls, the sustained proliferation of activated cells and their migration out of tissue niches will be driven by local cues, potentially wound-epidermis factors (**Figure 7B**). Understanding how wound epidermis-derived cues influence the biology of systemically-activated cells will be important. We showed that the systemic activation and priming effects are transient and disappear within several weeks. The mechanism whereby the effects are extinguished remains to be determined and defining the precise roles of epigenetics in this process requires more experimentation. There is precedence for non-immune cells, including epidermal stem cells, epigenetically encoding prior “memories” of traumatic injury and these memories sensitizing or otherwise influencing future injury responses.^39,40^ Beyond transcription factor binding site accessibility, post-transcriptional and post-translational regulation of transcription factor activity in systemic activation warrants investigation.

Systemic activation can have short-term benefits to the organism by priming other tissues for regeneration. The consequences of this system over an individual lifespan need to be evaluated. Repeated injury and chronic stimulation of this system may come at a cost. We previously demonstrated that even the axolotl’s limb regenerative abilities can be compromised by repeated amputation-regeneration cycles,^41^ as is also the case in at least one species of newt;^42^ hence, investigating how regenerative decline provoked by repeated injury interfaces with systemic activation will be interesting.

Our work suggests mammals have evolved or retained additional molecular controls that restrict natural regenerative ability. We highlight an important role for sympathetic innervation and adrenergic signaling in systemic responses to amputation, neither of which has been evaluated in mice. Future work in mice should evaluate these roles and evaluate their connection to circulating factors. These differences may reflect evolutionary differences in how regeneration responses are coordinated. Axolotls may prioritize regeneration over other physiological processes, such as growth or tissue maintenance, when necessary. This possible trade-off merits future investigation. Future investigations may also consider how the processes we identified may contribute to organismal resiliency in species that frequently survive intense predation, such as elephant seals that viciously fight during mating season^43^ and marine mammalian species, such as dolphins, attacked by cookiecutter sharks.^44^

Overall, we demonstrate that investigating the biology of systemic activation is an effective means to unravel the key pathways used to prime cells for limb regeneration. Our work predicts that activating the relevant cell types and directing them toward a regenerative response could be stimulated by harnessing adrenergic signaling in a mammalian setting, providing a major advancement toward the goal of therapeutic limb regeneration in the future. Our work also offers a new perspective from which to consider possible effects of amputation in humans outside of the injury site. Our findings may also provide important reference points for medical applications outside of the amputee population, for example, in the context of “polytrauma,” in which a patient suffers severe wounds in several different body parts.^45^

### Limitations of the study

It remains to be determined whether epigenetic changes we observed may persist beyond the priming period. While we focus here on roles for the sympathetic nervous system in systemic activation, roles for sensory and motor peripheral innervation should also be explored. Possible contributions of circulating epinephrine (as a function of the hypothalamus-pituitary-adrenal axis) remains to be determined.

## ACKNOWLEDGMENTS

This work was supported by The Richard and Susan Smith Family Odyssey Award (J.L.W.), the NIH New Innovator Award DP2HD087953 (J.L.W.), NICHD R01HD095494 (J.L.W.), NICHD R01HD115272 (J.L.W.), NSF-CAREER IOS-2145925 (J.L.W.), Harvard University Faculty of Arts and Sciences (J.L.W.), Harvard/MIT Joint Basic Program in Neuroscience (J.L.W. and I.M.C.), NSF IOS-2421118 (M.V.P.), Studienstiftung des deutschen Volkes (T.F.), NIH T32-AR080622 (R.H.), the Human Frontiers Science Program Long-term Postdoctoral Fellowship (A.M.S.), the Harvard Program for Research in Science and Engineering (R.T.K., S.H. and J.L.), the Harvard College Research Program (S.H. and A.Y.L.W.), and the Harvard Stem Cell Institute (A.Y.L.W.). We thank Dr. Randal Voss and Dr. Jeramiah Smith for pre-publication access to the axolotl reference genome UKY_AmexF1_1. We thank Dr. Elaine Fuchs’ lab for sharing reagents. We thank Samantha Payne, Solsa Cariba, Sydney Chambule, Fabiana Duarte, Juliana Babu, Jeramiah Smith, Nataliya Timoshevskaya, Tomás Gomes, and Adnan Abouelela for technical assistance. We thank Julia Thulander, Kara Thornton, Isaac Adatto, Brianna Blackmore, Lauryn Wilson, Hayden Graham, Damian Bernard, Nicholas Cardelia, Erin Anderson, Vicky Yan, Omenma Abengowe, Rui Qun Miao, Mak Famulari, Nicole Mejia, Madison Hurley, Gregory Gundberg, William Ye, Tyler Mourey for animal care. We thank Ya-Chieh Hsu, Mansi Srivastava, Matthew Warman, Marc Kirschner, Davie Van Vactor, Paola Arlotta, Rich Lee, Paul Garrity, Cliff Tabin, and members of the Whited, Chiu, Greer and Lehoczky labs for discussions and comments on the manuscript. We thank Yulia Swartz for sharing reagents and assisting in data interpretation, the Harvard Center for Biological Imaging (RRID:SCR_018673) for infrastructure and support, and the Bauer Core Facility at Harvard University for cell sorting and sequencing.

## AUTHOR CONTRIBUTIONS

D.P.-D, J.C.P, R.T. K., S.E.W., R.H., B.E., S.H., J.L., J.V.M, M.V.P., I.M.C., and J.L.W. designed the experiments; V.B. and N.F contributed tiger salamander data; D.P.-D. J.C.P., T.F., S.G.J., S.J.B., S.E.W., E.K., L.V.C., M.V.P., I.M.C., and J.L.W. wrote the paper; T.F. S.G.J., S.J.B., H.S., E.K., S.Y.C.W., A.H.F., S.B., P.B., B.J.H, and T.B.S. designed and performed computational analyses; L.V.C. performed statistical analyses; D. P.-D., J.C.P, H.S., R.T. K., S.E.W., R.H., A.M.S., B.E., N.L., S.M., A.E.S., B.G., K.E.D., S.H., J.V.M., A.W., B. T., G. K., E. W. and J.L. performed sample preparation, experiments, and experimental data analyses. All authors contributed to manuscript editing.

## STAR Methods

### RESOURCE AVAILABILITY

#### Lead contact

Further information and requests for resources and reagents should be directed to and will be fulfilled by the lead contact, Jessica Whited.

#### Materials availability

This study did not generate new unique reagents.

#### Data and code availability

- All single-cell RNA-sequencing datasets generated in this study are available at Gene Expression Omnibus (GEO), accession numbers GSE232083 (4C) and GSE232082 (2C).
- Bulk ATAC-seq dataset generated in this study is available at Gene Expression Omnibus (GEO), accession number GSE232080.
- All original code has been deposited at GitHub and is publicly available as of the date of publication. The full URL is available in the Key Resources Table.
- Any additional information required to reanalyze the data reported in this paper is deposited at the Harvard Dataverse and available from the lead contact upon request.

### EXPERIMENTAL MODEL AND SUBJECT DETAILS

#### Animals

All animal experimentation was approved by and conducted in accordance with Harvard University’s Institutional Animal Care and Use Committee. Leucistic axolotls were used for all animal experiments except RNA sequencing and maintained as previously described.^29^ For RNA sequencing, *βIIItub:GAP43-EGFP* transgenic animals were used.^66^ Animals in all experiments were both age and size matched. Animals were anesthetized in 0.1% (w/v) tricaine prior to all procedures involving amputation, denervation, or injection. After all surgical procedures, axolotls were allowed to recover overnight in 0.5% (w/v) sulfamerazine. All limb amputations were performed at the mid-stylopod and the bone was then trimmed back from the amputation plane. For forelimb denervations the brachial plexus nerves were transected as described previously,^67^ and for hindlimb denervations the sciatic nerves were transected as described^68^. Sham operations as controls for denervations were performed by anesthetizing animals and cutting the skin open with sterile scissors, locating the nerve bundles with closed sterile forceps, but not cutting the nerves.

### METHOD DETAILS

#### Denervations and amputations

Animals in all experiments were both age and size matched. Animals were anesthetized in 0.1% (w/v) tricaine prior to all procedures involving amputation, denervation, or injection. After all surgical procedures, axolotls were allowed to recover overnight in 0.5% (w/v) sulfamerazine. All limb amputations were performed at the mid-stylopod and the bone was then trimmed back from the amputation plane. For forelimb denervations the brachial plexus nerves were transected according to Schotté *et al.* ^67^ and for hindlimb denervations the sciatic nerves were transected according to Kropf *et al.* ^68^ Sham operations as controls for denervations were performed by anesthetizing animals and cutting the skin open with sterile scissors, locating the nerve bundles with closed sterile forceps, but not cutting the nerves.

#### Drug treatments and EdU incorporation

Rapamycin was dissolved into a 25 mg/mL stock solution in ethanol. Rapamycin treatments were delivered via both water treatments at a concentration of 1 µM for 16 days with water changed daily, and through daily intraperitoneal injections at a concentration of 5 mg/kg/day for 16 days, both starting two days before amputation. All intraperitoneal injections were administered at a volume of 20 µL/g in a vehicle solution comprised of APBS with 5% Tween80, 5% PEG-400. All stock solutions were frozen at −80 °C in 1-mL aliquots. A stock solution of 6-hydroxydopamine hydrochloride (Sigma-Aldrich H438) was prepared in 0.01% ascorbic acid at a concentration of 15 mg/mL and was administered intraperitoneally for two consecutive days at a dose of 300 mg/kg. The animals were allowed to recover for 5 days before amputations. Yohimbine (Sigma-Aldrich Y3125) was dissolved into a 10 mg/mL stock solution with sterilized water. Yohimbine was administered daily at a dose of 1mg/kg for 14 days via intraperitoneal injection. Clonidine (Sigma-Aldrich C7897) was dissolved in water at 7.5 mM stock concentration and was delivered via water treatments at 30 µM. A stock solution of propranolol (Sigma-Aldrich P0884) was created by dissolving 100 mM in water. Propranolol was delivered via water treatments at 5 µM. 5-ethynyl-2′-deoxyuridine (EdU) was purchased from ThermoFisher and stock solutions were created by dissolving the powder in dimethyl sulfoxide as instructed by the manufacturer. Following anesthetization, axolotls were administered 20 µL/g of 400 µM EdU in 0.7X PBS via intraperitoneal injection 18 hours before tissue harvest.

#### Skeletal and tissue preparations

For immunofluorescence and HCR-FISH experiments, tissue samples were fixed in 4% paraformaldehyde post-collection in DEPC-treated PBS overnight at 4°C. Tissues were then cryopreserved in a sucrose gradient following a wash in DEPC-PBS and incubated in 30% sucrose/DEPC-PBS at 4°C overnight. Tissues were then embedded and frozen on dry ice in Tissue-Tek O.C.T. Compound (Sakura) and stored at −80°C. All blocks were sectioned at 16 μm thickness using a Leica CM 1950 cryotome and sections were stored at −80°C until use.

For wholemount RNAscope, forelimbs were fixed in 4% paraformaldehyde at 4°C overnight. Limbs were dehydrated with 50% methanol, 70% methanol, and 100% methanol. Dehydrated limbs were pre-treated with RNAscope target retrieval (ACD 322000) and protease plus (ACD 322331).

For skeletal preparations, post-collection, limbs were incubated overnight in 95% ethanol at room temperature and then incubated in 100% acetone under the same conditions. Skeletal elements were visualized with Alcian blue and Alizarin red S stains followed by a clearing of non-skeletal tissue with potassium hydroxide.

#### Immunofluorescence, HCR-FISH and RNAscope

Sections were rehydrated with PBS and then permeabilized with 0.5% Triton-X-100 PBS. Post-permeabilization, slides were rinsed twice with PBS and EdU visualization was performed via sulfo-cyanine 3 azide (Lumiprobe). Post-EdU visualization slides were washed twice with PBS. Staining and labeling with any additional antibodies was conducted post-EdU labeling. Primary antibodies used were rabbit anti-phospho-S6 ribosomal protein (Ser235/236, 1:200; Cell Signaling 4858), rabbit anti-tyrosine hydroxylase (1:200, Invitrogen PA5-85167). Secondary antibodies used were Cy3-goat anti-mouse (1:250, Jackson ImmunoResearch 115-165-146), Alexa Fluor 488-Goat Anti-Rabbit (1:250, Jackson ImmunoResearch, 111-545-003) and Cy3-goat anti-rabbit (1:250, Jackson ImmunoResearch 111-165-003).

Sections stained with all antibodies were blocked with 10% serum from the host species of the secondary antibody in 0.1% Triton-X-100 in PBS. Primary and secondary antibodies were diluted in in 2% BSA, 0.1% Triton-X-100 in PBS. Sections labeled with anti-pS6 and NeuN underwent an antigen retrieval step after permeabilization, in which slides were boiled in 0.1 M sodium citrate (pH 6.0) and then rinsed once in PBS, before EdU was visualized, followed by blocking and primary antibody incubation. All slides were incubated with DAPI (Sigma D9542) before cover slipping.

HCR probes were designed using the Probe Generator tool (Monaghan Lab, Northeastern University, MA, USA). HCR-FISH was performed as described previously.^69^ For RNAScope, probes were hybridized and fluorescently labeled with RNAscope Multiplex Fluorescent Detection Reagents kit (323110, ACD) following the manufacturer’s instruction. After, limbs were incubated in 1.3g/mL Histodenz (Sigma-Aldrich D2158) in PBS for up to 3 days. Samples were imaged using Zeiss lightsheet Z.1. Images were analyzed using Imaris software (Bitplane).

#### Cell Culture

AL-1 cells are derived from axolotl limb dermal fibroblasts^16,70^. AL-1 cells were maintained as described in Lévesque et al.^71^. Cells were kept at 26°C without CO_2_. They were grown in complete AL-1 media: 60% L-15 (Sigma Aldrich L5520), 5% Fetal Bovine Serum (FBS) (VWR 1500–500), 200 U/mL penicillin and 200 µg/mL streptomycin (Thermo 15-140–122), 250 ng/mL amphotericin B (Thermo 15240062), 2 mM L-Glutamine (Thermo 25-030–081), 10 µg/mL Insulin, 5.5 µg/mL transferrin, and 6.7 ng/mL sodium selenite (Thermo 41400045).

#### Cell Line Drug Treatments and Immunofluorescence

Cells were seeded at 100,000 cells per well in 12-well glass bottom plates. Cells were given 100 µM epinephrine diluted in water, 200nM rapamycin diluted in DMSO, vehicle control, or both 100 µM epinephrine and 200nM rapamycin. After 24 hours of treatment, cells were washed in PBS, fixed in 4% PFA for 10 minutes, rinsed in PBS, and permeabilized with 0.5% Triton-X-100 PBS. After permeabilization, cells underwent antigen retrieval where samples were boiled in 0.1 M sodium citrate (pH 6.0) and rinsed in PBS before subsequent blocking in 10% donkey serum in 0.5% Triton-X-100 PBS, overnight primary antibody incubation, secondary antibody incubation, and DAPI staining (Sigma D9542). Primary antibody used was rabbit anti-phospho-S6 ribosomal protein (Ser235/236, 1:200; Cell Signaling Technology 4858) and secondary antibody used was Alexa Fluor 488-Goat Anti-Rabbit (1:250, Jackson ImmunoResearch 111-545-003).

#### Imaging and quantification

Imaging was performed on a Zeiss Axio Scan.Z1 automated slide scanner, Leica M165 FC equipped with Leica DFC310 FX camera, a Nikon Eclipse Ni microscope with a DS-Ri2 camera, and a Nikon Spinning Disk Confocal (CSU-W1). Quantification of all images was conducted blindly. For imaging of homeostatic (intact), contralateral or regenerating limbs, representative sections displaying all non-epidermal tissue types were selected. For the tissue type analysis of EdU+ nuclei, tissues were classified as epidermis, “skeletal elements” (bone, cartilage), and “soft tissues” (muscle, joint, tendon, ligaments, dermis, vasculature, and nerves) based on nuclear morphology and location as previously described (Whited et al, 2013). All cell counts were conducted utilizing CellProfiler.^72^ Measurements of blastema area were performed using Fiji.^48^ For HCR signal quantifications, the wound epidermis was first masked from each image. HCR signal’s fluorescence intensity was normalized to the area of the blastema as measured by the DAPI signal.

#### Cell dissociation, FACS isolation, and single-cell RNA-sequencing

To isolate single cells, 12-16 homeostatic or systemically-activated limbs at 14 days post-amputation were pooled for each biological replicate and dissociated in 4 ml of dissociation buffer (100 mg/ml liberase diluted in 0.7X PBS with 0.5 U/ml DNase I) for 30 min at room temperature with gentle mixing. Suspensions were filtered through 70 µm strainers to remove cell clumps and debris, and strainers were washed with 1 ml of 0.7X PBS. Cells were spun down at 300 g at room temperature and resuspended in 1 ml of 0.7X PBS-0.04% BSA. Cells were then stained with 1 mg/mL Hoecsht, calcein AM, and propidium iodide for isolation of alive cells. Cells were further sorted into dividing (4C) and non-dividing (2C) cells by fluorescence activated cell sorting. 4C and 2C cell fractions originating from different pools of limbs were purified and were then sequenced at the Bauer Core Facility at Harvard University using an Illumina NovaSeq 6000 [https://www.illumina.com]. Paired end reads were demultiplexed with bcl2fastq2 (V2.2.1) (RRRID:SCR_015058).

#### ATAC-seq Library Prep and Sequencing

Ten thousand sorted 4C cells from homeostatic, systemically-activated, and regenerating limbs were resuspended in 5 μL PBS. 42.5 μL of transposition buffer (38.8 mM tris-acetate, 77.6 mM potassium acetate, 11.8 mM magnesium acetate, 0.12% NP-40, 18.8% DMF, 0.47% protease inhibitor cocktail) was added to the cells, mixed, and incubated for 10 min at room temperature. 2.5 μL of pre-loaded Tn5 (Diagenode C01070012) was added to the reaction mixture. The transposition reaction was carried out at 37°C for 30 min with shaking at 300 rpm. Following transposition, the samples were purified through DNA Clean and Concentrator-5 Kit (Zymo Research D4014). Purified library fragments were amplified with custom primers 1 and 2 using the following conditions: 72°C for 5 min; 98°C for 30 s; and thermocycling at 98°C for 10 s, 72°C for 30 s and 72°C for 1 min. The libraries were amplified for five cycles. Soon after, 5 μL aliquot of the PCR reaction was added with 10 μL of the PCR cocktail with SYBR Green (0.6X final concentration). 20 cycles of PCR was performed to determine the additional number of cycles needed for the remaining 45 μL reaction. The libraries were purified using DNA Clean and Concentrator-5 Kit (Zymo Research D4014). Purified libraries were sequenced at The Bauer Core Facility at Harvard University using an Illumina NovaSeq 6000 [https://www.illumina.com]. Paired End reads were demultiplexed with bcl2fastq2 (V2.2.1) (RRID: SCR_015058).

### QUANTIFICATION AND STATISTICAL ANALYSIS

#### Single-cell RNA-sequencing read quantification

2C and 4C datasets were preprocessed independently and only merged after all quality control steps. To quantify reads, python v3.9.21 and the kb python package (v0.29.1) ^73^ was used. Within this workflow, kallisto (v0.51.1) ^49^ was used to create a reference index and to pseudoalign reads. Bustools (v0.44.1) ^50^ was used to quantify the output. For all steps, the latest axolotl genome assembly UKY_AmexF1_1 was used (Accession number: JBEBLI000000000.1, RefSeq assembly: GCF_040938575.1) (https://ambystoma.uky.edu/genome-resources). Matrix files were then read into R Studio (v 4.3.3) and a Seurat object (v5.0.3) ^51^ was created. Empty droplets were filtered out using the emptyDrops command within DropletUtils (v1.22.0) ^52^, where an FDR >0.001 was chosen as a cutoff value. Further, cells with <500 UMIs were removed, to only include cells with adequate sequencing depth. Data was clustered preliminarily in Seurat as input to SoupX (1.6.2) ^53^ to correct for ambient RNA contamination. SoupAdjusted counts were used to filter out doublets via the scDblFinder package ^54^ (v1.16.0). Finally, dying cells (>5% mitochondrial gene counts) were removed.

2C and 4C datasets were independently normalized on their respective SoupAdjusted layer using the *SCTransform* command ^74^, and potential mitochondrial mapping confounding variables were accounted for via the *vars.to.regress* parameter. 2C and 4C datasets were finally merged and integrated using Harmony ^55^ (v1.2.0). Cells were clustered in Seurat with 15 dimensions and at a resolution of 0.15 and visualized with the UMAP reduction method. Marker genes were determined via thze *FindAllMarkers* command, where only genes expressed in >20% of cells in a given cluster, with an associated log2 fold-change >0.25 were considered. These marker genes were used to annotate clusters manually.

To determine co-expression of *Adra2a* with blastema marker genes, cells were categorized into four groups. If cells expressed >0 counts of *Adra2a*, they were flagged as Adra2a+, while all other cells were classified as Adra2a-. Cells were further divided into homeostatic and activated, dependent on their experimental condition. Next, the Seurat command *FindAllMarkers* was deployed to test whether common blastema marker genes ^75^ were differentially expressed, with a significance-threshold of 0.01. Data was further visualized via the *DotPlot* command within Seurat.

### ATAC-seq read processing

We aligned sequencing reads to the reference genome (AMEX60DD) using Bowtie2 (v2.3.2)^56^ with parameters optimized for local alignment (--no-mixed), high sensitivity (-X 2000), gapped alignment (--dovetail), and exclusion of discordant read pairs (--no-discordant). Alignments were sorted and indexed throughout the analysis using Samtools (v1.10)^57^ for efficient retrieval. Mitochondrial reads were filtered out using grep to focus on nuclear DNA.Low-quality alignments (MAPQ score < 30) were removed using Samtools. PCR and optical duplicates were identified and removed with Picard MarkDuplicates^58^ [https://broadinstitute.github.io/picard/] assuming sorted alignments (ASSUME_SORTED=true) and removing all identified duplicates (REMOVE_DUPLICATES=true). Generation of bigwig Coverage Tracks was performed with deepTools bamCoverage^76^. We specified ––outFileFormat=bigwig for bigWig output, –– normalizeUsing CPM for counts per million normalization, and ––binSize 20 for a bin size of 20 base pairs.

### ATAC-seq peak calling

Open chromatin regions were identified using MACS2 (v2.2.9.1)^59^ (https://github.com/macs3-project/MACS). We employed two peak calling strategies: consensus peaks, generated by analyzing all replicates/samples together, and condition-specific peaks.

### Annotating and visualizing the ATAC data

We employed ChIPseeker (v1.32.1)^60^ to annotate the peaks. Subread featurecount (v1.32.1)^77^ was used to create a gene by peak count matrix. Differential accessibility was observed using DESeq2.^61^ To visualize normalized counts and sample clustering, we used Pheatmap (v1.9.12) (<u>10.32614/CRAN.package.pheatmap</u>) ^62^ and pcaExplorer (v2.22.0)^63^, respectively. Motif discovery and enrichment analysis were performed using HOMER (v4.11).^64^ Briefly, motifs from HOMER analysis were read into MATLAB and significant motifs identified in each replicate for each condition. Replicates with number of enriched motifs significantly lower than in other replicates for the same condition were discarded in further analyses. 4C single-cell gene expression linked to discovered motifs was visualized with Seurat’s DotPlot function.^65^ We preprocessed the 4C data using Seurat’s AggregateExpression function with log normalization and scaling. Heatmaps of the aggregated 4C single cell counts were generated with pheatmap (<u>10.32614/CRAN.package.pheatmap</u>)^62^. Finally, pathway enrichment analysis of the transcription factors was performed using Panther^78^ and visualized with ggplot2.^79^

### SUPPLEMENTAL FIGURE LEGENDS

**Figure S1.**
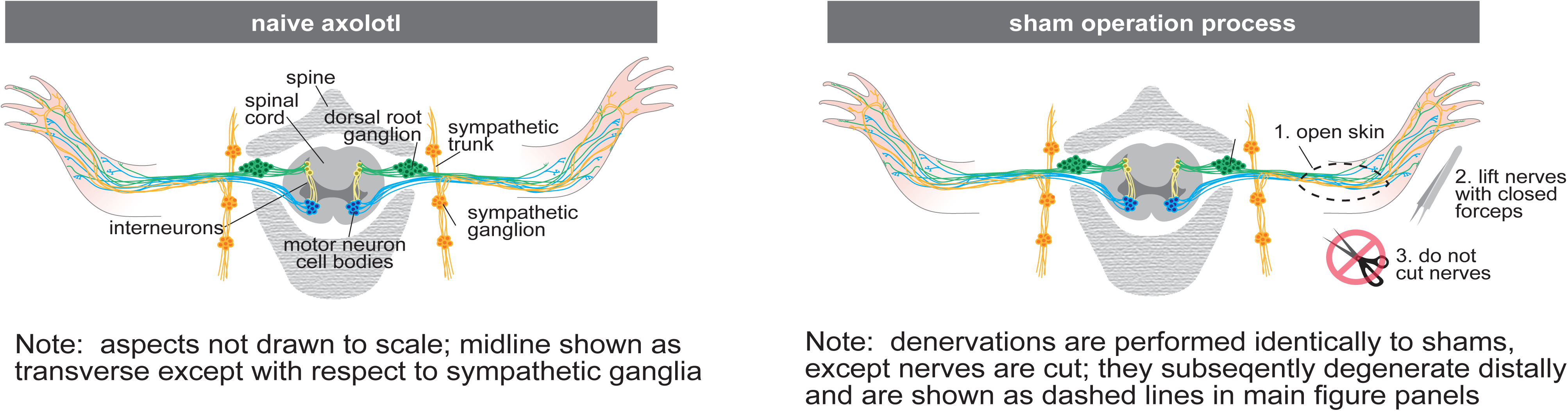
Denervation surgery (Related to Figure 2). Cartoon depicting nerves innervating the axolotl limb in the naïve state (left) and the sham-operation procedure (right).

**Figure S2.**
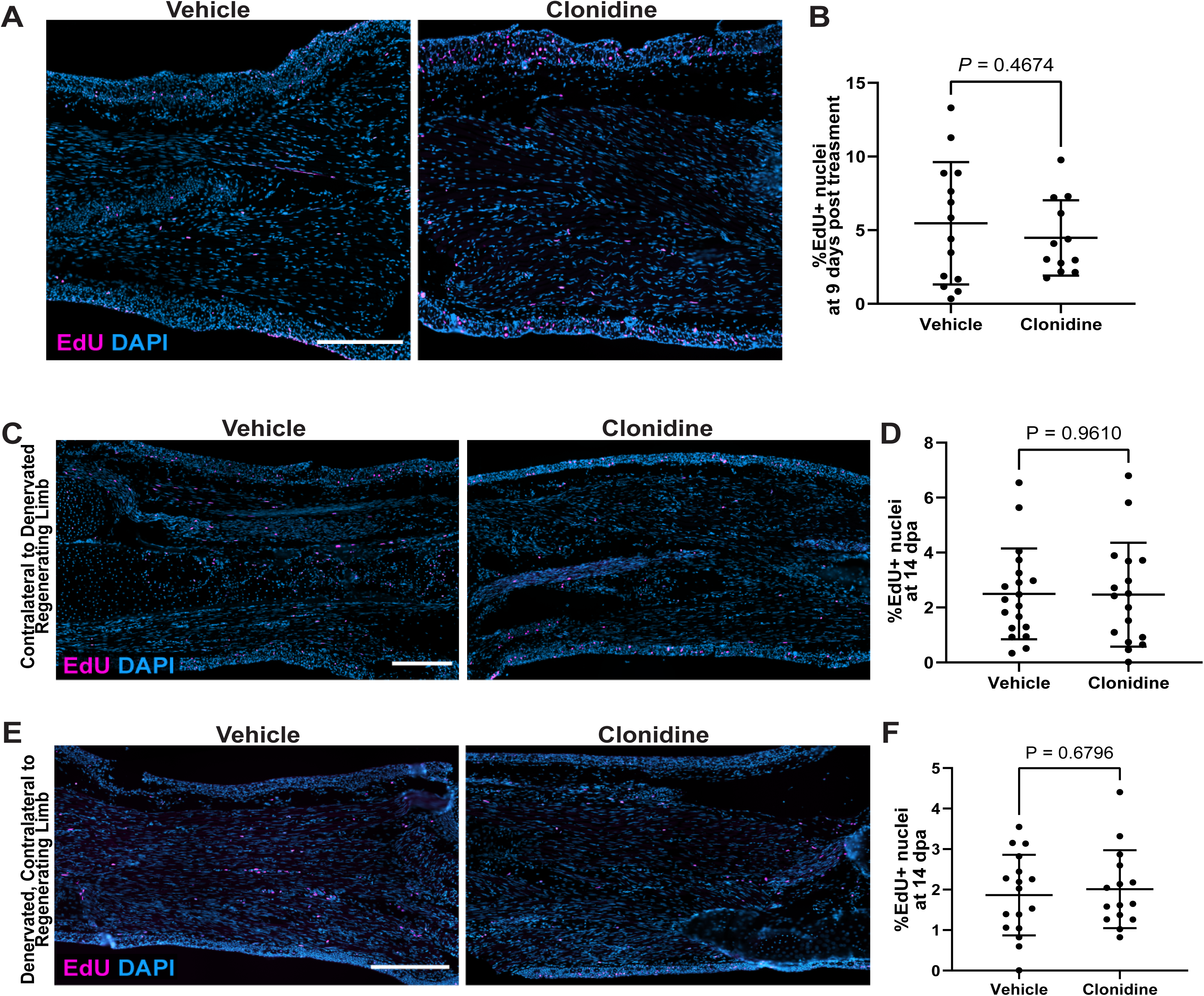
Activation of α-adrenergic signaling is not sufficient for systemic activation (Related to Figure 3). **(A)** Representative images of EdU and DAPI stained intact limbs harvested from animals treated with Adra2a agonist clonidine or vehicle solution for 9 days. **(B)** Quantification of percent EdU positive nuclei in the intact limb (n = 14 animals for vehicle and n = 12 for clonidine). **(C-D)** Animals treated with Adra2a agonist clonidine or vehicle solution were unilaterally denervated followed by proximal amputations. **(C)** Representative images of EdU and DAPI stained limbs contralateral to denervated and amputated limbs, harvested at 14 days post-amputation. **(D)** Quantification of percent EdU positive nuclei of contralateral limbs in **(C)** (n = 19 animals for vehicle and n = 17 for clonidine). **(E-F)** Animals treated with Adra2a agonist clonidine or vehicle solution were unilaterally denervated on one side, followed by unilateral proximal amputations of the innervated limbs. **(E)** Representative images of EdU and DAPI stained limbs, denervated and contralateral to unilateral amputations, harvested at 14 days post-amputation. **(F)** Quantification of percent EdU positive nuclei of denervated contralateral limbs in **(E)** (n = 17 animals for vehicle and n = 15 for clonidine). Statistical significance was determined using unpaired two-tailed Welch’s t-test in **(D)** and **(F)**. Scale bars, 500 µm. EdU, 5-ethynyl-2′-deoxyuridine.

**Figure S3.**
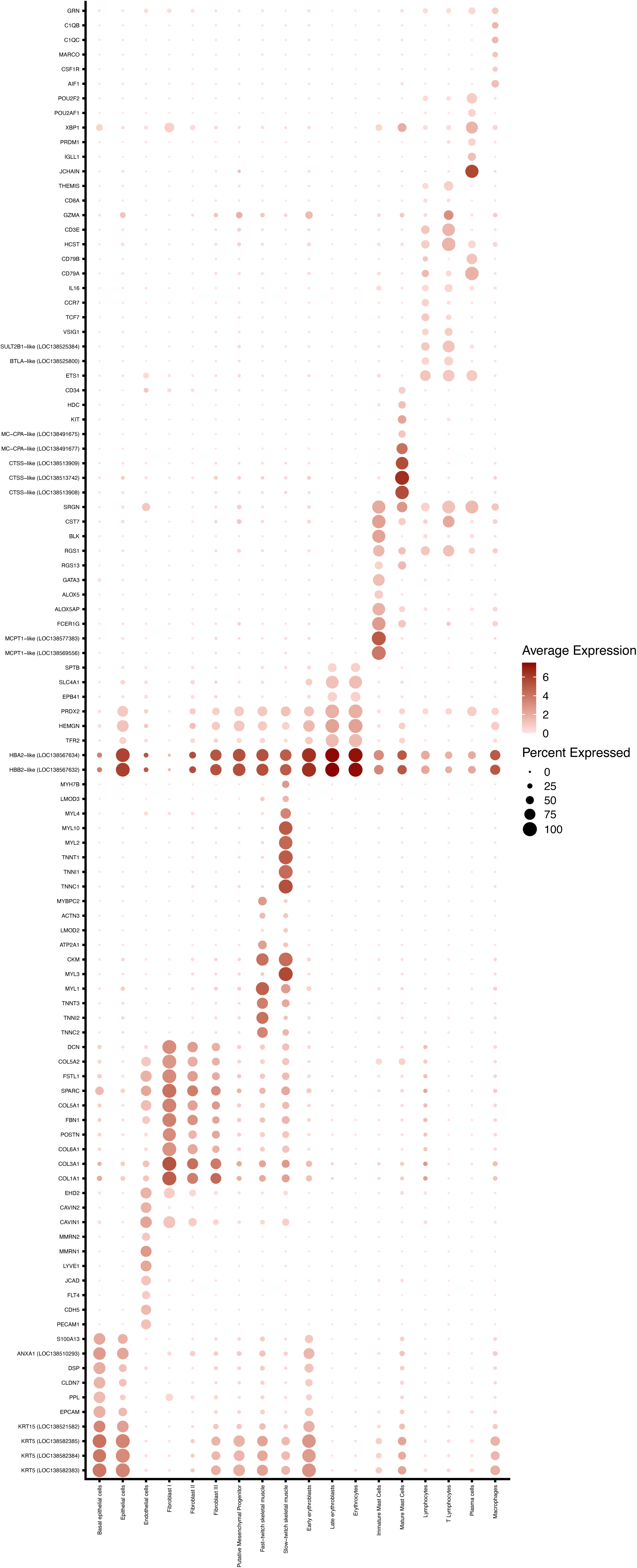
Marker genes used to identify clusters in single-cell RNA sequencing of 4C and 2C cells (Related to Figure 6). Dot plots showing the relative frequency of expression (dot size) and expression level (color) of marker genes across all clusters.

**Figure S4.**
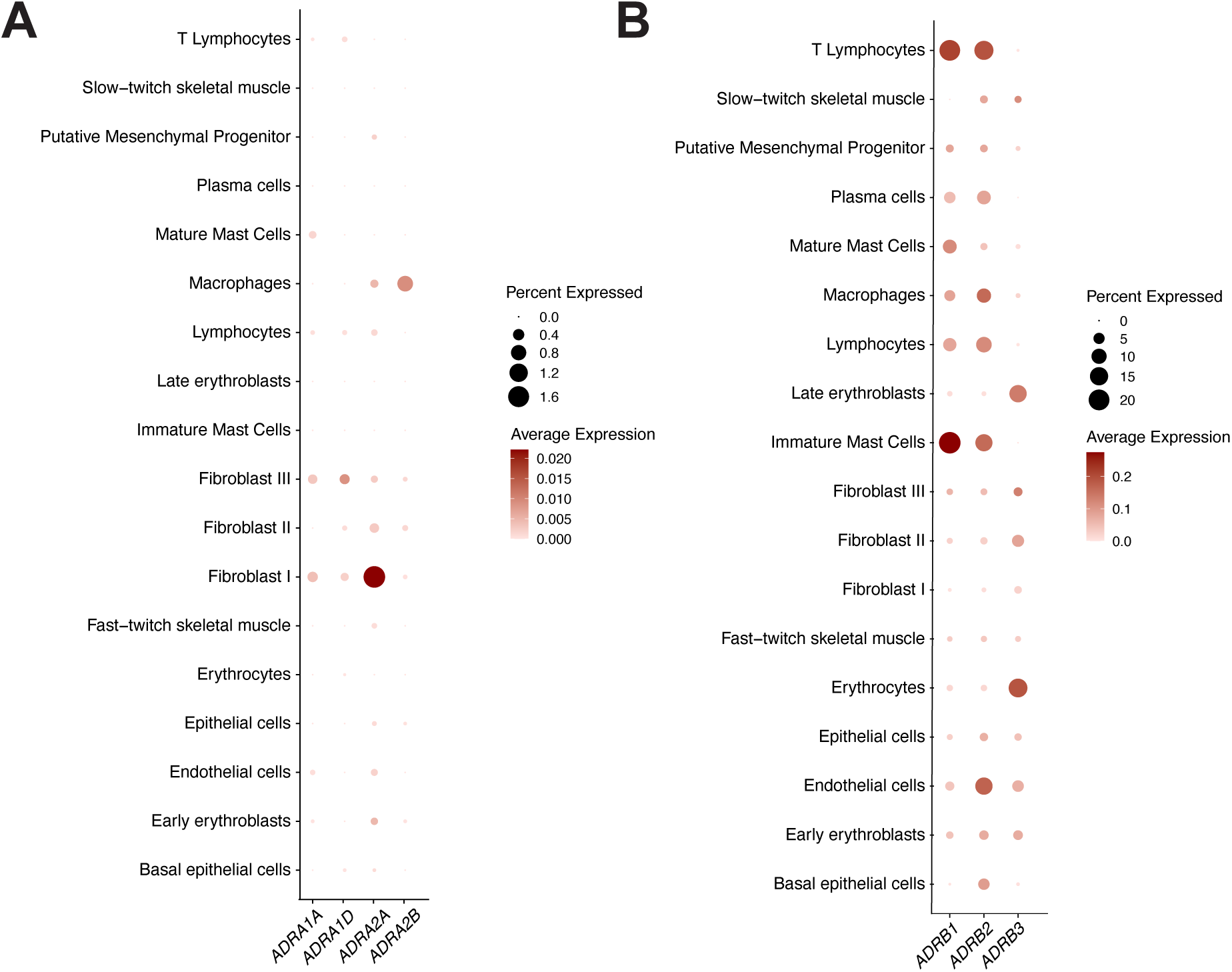
Expression of adrenergic receptors across cell types (Related to Figure 6). **(A-B)** Dot plots showing the relative frequency of expression (dot size) and expression level (color) for alpha-**(A)** and beta-**(B)** adrenergic receptors across all cell clusters.

**Figure S5.**
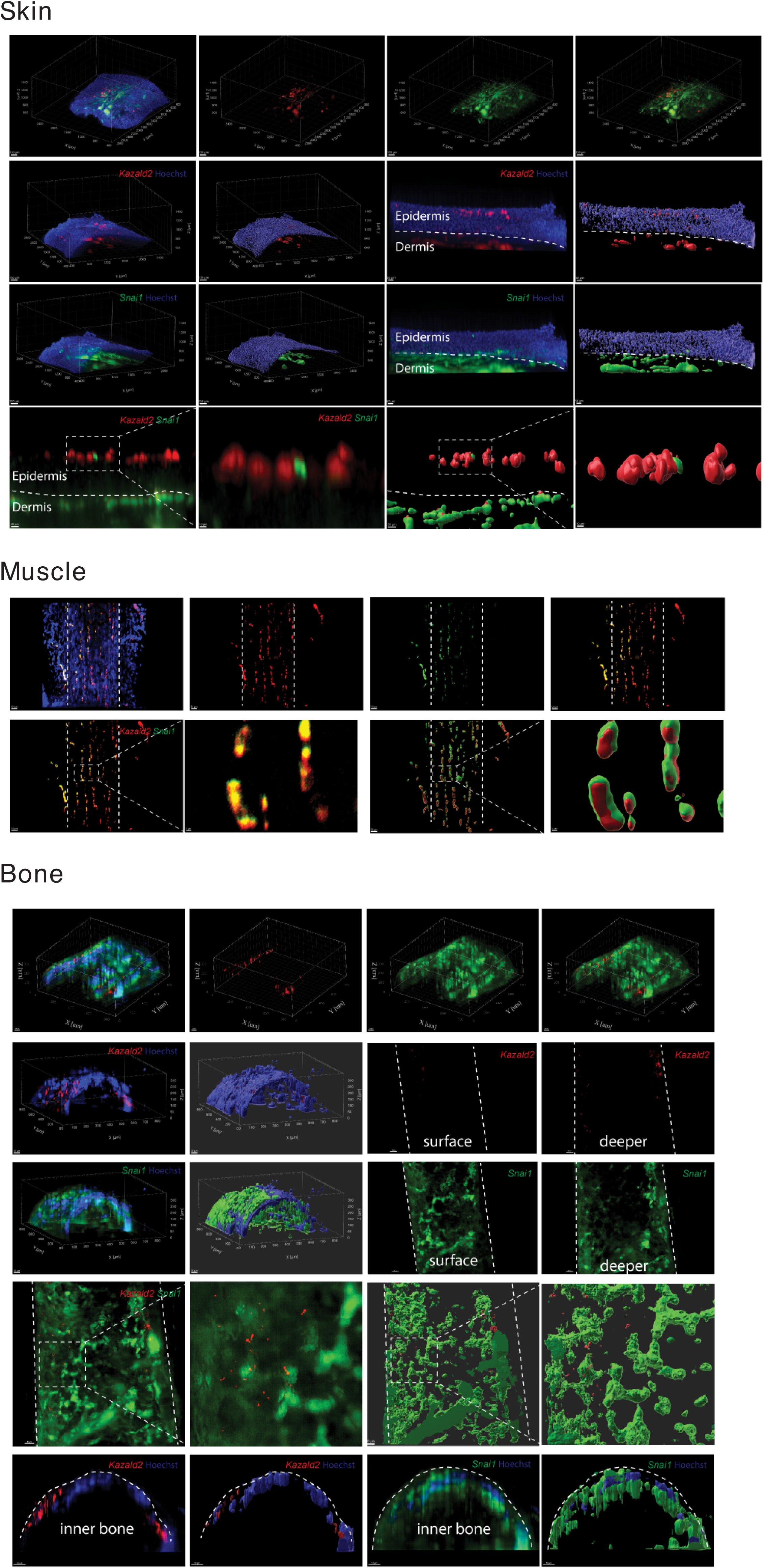
Expression of *Kazald2* and *Snai1* in the wholemount homeostatic limb (Related to Figure 6). Representative images of wholemount limbs showing *Kazald2* and *Snai1* expression in the skin (top panel), muscle (middle panel) and bone (bottom panel).

### SUPPLEMENTAL EXCEL TABLE LEGENDS

**Table S1.** List of transcripts expressed by each cell cluster in homeostatic and systemically-activated limbs (Related to Figure 6).

**Table S2.** Differentially expressed transcripts expressed by each cell cluster in homeostatic and systemically-activated limbs (Related to Figure 6).

**Table S3.** List of accessible transcription factor binding motifs enriched across 4C cells of homeostatic, activated, and regenerating limbs. (Related to Figure 6).

### SUPPLEMENTAL MOVIE LEGENDS

**Supplementary Movie 1. Visualization of whole mount bone of a homeostatic axolotl limb showing *Kazald2* (red) and *Snai1* (green) expression (Related to Figure 6).**

---

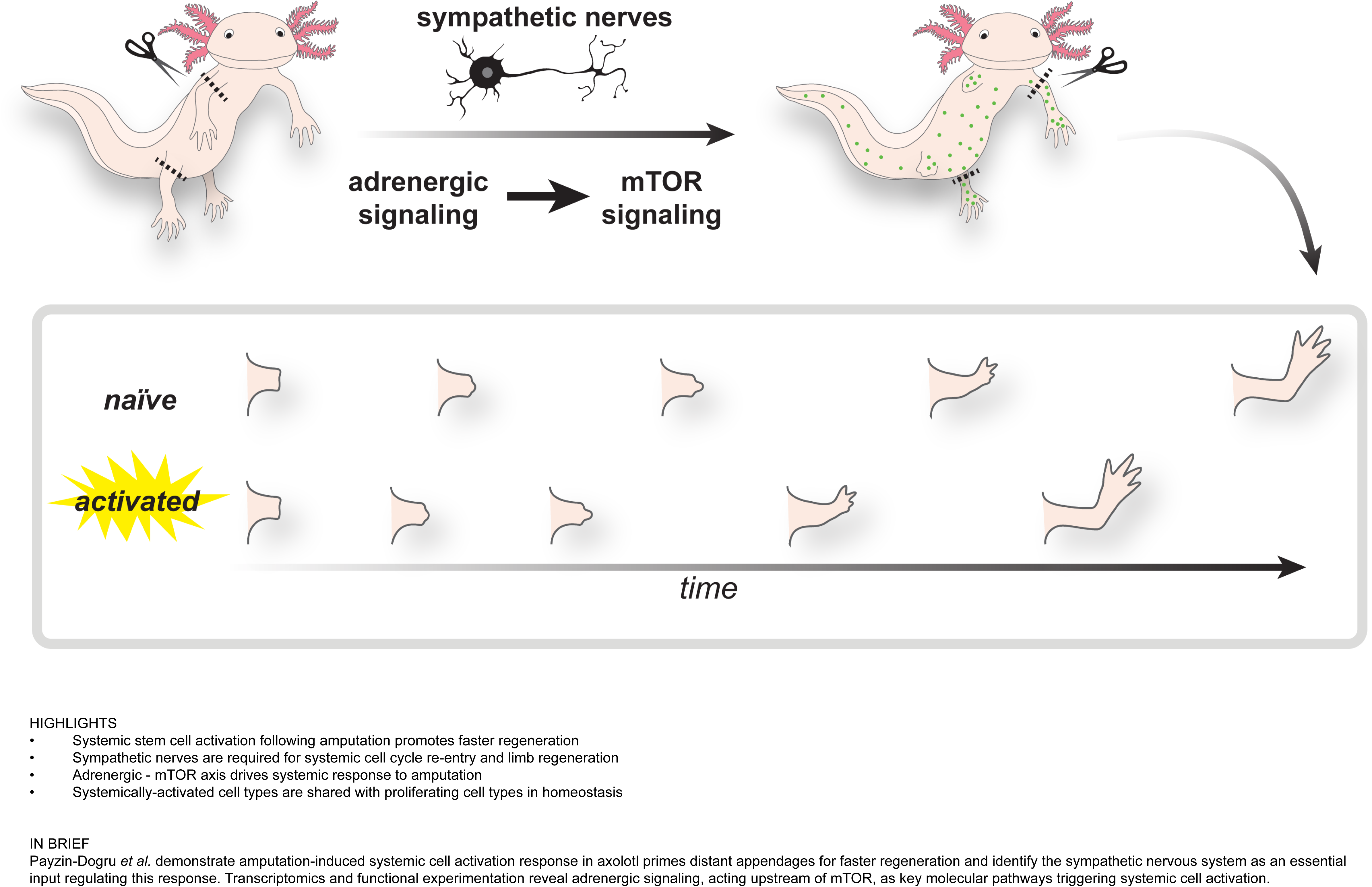

